# Characterization and Bioactivity Potential of Marine Sponges (*Biemna fistulosa, Callyspongia diffusa,* and *Haliclona fascigera*) from Kenyan Coastal Waters

**DOI:** 10.1101/2025.05.20.655062

**Authors:** Teresia Nyambura Wacira, Huxley Mae Makonde, Joseph Kamau Nyingi, Christopher Mulanda Aura, Cromwell Mwiti Kibiti

**Author notes:** Corresponding Author: Teresia Nyambura Wacira. Huxley M. Makonde, Joseph K. Nyingi,8 Christopher M. Aura, Cromwell M. Kibiti.

## Abstract

Sponges have been reported as a rich source of bioactive compounds, which could potentially be developed into lead compounds for pharmaceutical use. The present study aimed to identify sponges and assess the biological activity of their extracts against human disease-causing organisms, including *Escherichia coli*, *Pseudomonas aeruginosa, Staphylococcus aureus,* and *Candida albicans*. Morphological characterization and DNA barcoding of the cytochrome c oxidase subunit I (COI) gene characterized three sponge species (*Biemna fistulosa*, *Callyspongia diffusa* and *Haliclona fascigera*). The Kirby-Bauer test assessed the antimicrobial activity of the extracts and the inhibition zone diameters (IZD) were measured. The extracts were further subjected to minimum inhibitory concentration (MIC) and minimum bactericidal concentration (MBC) tests to determine the antibiotic susceptibility. The Gas Chromatography-Mass Spectrometry (GC-MS) was used to identify and quantify the organic compounds in the sponges’ extracts. The methanolic extract of *B. fistulosa* (28.00 ± 3.5 mm) and *H. fascigera*. (28.33 ± 3.8 mm) exhibited a broad spectrum of antibacterial activity against *E. coli* surpassing the positive control (27.67 ± 0.9 mm). The inhibitory activity of ethyl acetate extract of the *C. diffusa* (29.33 ± 2.4 mm) against *P. aeruginosa* was observed to be higher compared to the standard antibiotic streptomycin (26.67±0.7 mm). The methanolic extract of *H. fascigera* demonstrated the lowest MIC (0.53 ± 0.0 mg/ml) in compared to the streptomycin drug (1.36 ± 0.0 mg/ml), and showed an MBC of 1.25 mg/ml against *E. coli*. The GC-MS chromatogram data analysis identified 114 distinct compounds categorized into 39 classes across three sponge extracts, with 11.4% of these compounds demonstrating documented antimicrobial activity against human pathogens. This study corroborates sponges as a promising source of bioactive compounds, which are valuable leads for drug discovery and development. Future research must explore their mechanisms, molecular-level toxicity, and lead optimization to enhance drug development.

## Introduction

Despite a record dating back at least the entire Phanerozoic, approximately 600 million years (Carrier et al., 2022). The magnitude of aquatic marine sponge biodiversity is unknown (Folkers and Rombouts, 2020). As of March 2025, the World Porifera Database (WPD) (Soest et al., 2019) recognizes approximately 9,722 accepted living sponge species and 60 extinct species. In 1950, Bergmann and Feeney discovered marine sponge natural products by extracting novel spongothymidine and spongouridine nucleosides from *Tectitethya crypta*, formerly known as *Cryptotheca crypta* (Kurian and Elumalai, 2017). Among the three classes of sponges, class Demospongiae and orders Halichondrida, Poecilosclerida, and Dictyoceratida are the significant sources of bioactive compounds compared to Hexactinellida and Calcarea (Sharma et al., 2010). Recent scholarly investigations underscore the considerable potential of marine sponges as a source for novel antimicrobial agents. Notably, a study conducted on sponges from Saint Thomas, U.S. Virgin Islands, demonstrated significant antibacterial activity against several pathogenic microorganisms, including *Escherichia coli, Staphylococcus aureus*, and *Neisseria gonorrhoeae* (Christian et al., 2024). More than 200 new compounds have been isolated from marine sponges, accounting for approximately 23% of approved marine-derived pharmaceuticals (Hong et al., 2022).

Dragmacidin G, a bioactive alkaloid extracted from sponges of the genera *Spongosorites* and *Lipastrotheya*, exhibited a broad spectrum of biological activity, including the inhibition of *Mycobacterium tuberculosis*, *Plasmodium falciparum* and methicillin-resistant *Staphylococcus aureus* (MRSA) (Wright et al., 2017). Manzamine A, derived from Indo-Pacific sponges, manifests broad-spectrum antibacterial activity against both Gram-positive and Gram-negative bacterial strains (Santi et al., 2025). Similarly, Aeroplysinin-1, extracted from *Aplysina aerophoba*, has revealed significant antibacterial properties against various pathogenic strains (García-Vilas et al., 2016). Additionally, compounds such as Discorhabdin G and Ageliferin from sponges have been recognized for their ability to prevent bacterial biofilm formation, a critical factor that contributes to the persistence and resilience of infections (Huigens, 2009). Furthermore, the pharmacological potential of marine sponges transcends antibacterial activity, as they are also abundant sources of antifungal and antiviral agents. Avarol, a hydroquinone derivative isolated from the sponge *Dysidea avara*, has antifungal activity against various pathogenic fungi (Esposito et al., 2022). Mycalamide A, obtained from *Mycale* sp., is renowned for its antiviral activity, particularly against herpes simplex virus and poliovirus type 1 (Sharma et al., 2016).

Despite the significant advancements made in the research of marine sponges for the production of bioactive compounds, considerable gaps remain in our understanding of marine sponges and their bioactive compounds (Mehbub et al., 2024). Furthermore, while various sponge-derived compounds are recognized for their antibacterial and antifungal activities, there is inadequate literature and comprehensive data on the identification and bioactivity studies of marine sponges from the Kenyan coastal waters. (Handayani et al., 2023). This scientific gap impedes the progression of sponge bioactive compounds from laboratory studies to clinical applications, a situation exacerbated by the scarcity of high-throughput screening technologies (Ayon, 2023). Numerous coastal and deep-sea sponge species, particularly those inhabiting African waters, such as those off the coast of Kenya, remain largely underexplored (Garritano et al., 2023). To bridge the scientific gap, the present comprehensive study identified some marine sponge species and characterized their antimicrobial potential against selected human pathogens.

## MATERIALS AND METHODS

### Description of study site

The study sites included Sii Island (4°40’46.1"S, 39°17’01.9"E), Mundini (4°39’10.0"S, 39°21’41.0" E), and Ras Kiromo (4°38’45.5"S, 39°19’23.4"E), located on the South Coastline of Kenya. These are sites characterized by diverse marine ecosystems, with coral reefs and seagrass beds supporting a wide range of aquatic organisms (Musembi et al., 2019).

### Metazoan specimen collection and preparation

A pre-survey for marine sponge sampling was undertaken to determine optimal conditions for clear underwater photography and effective sampling during favorable weather conditions (Cai et al., 2024). The collection of metazoan marine sponges took place during the low spring tide, coinciding with the southeast monsoon in September 2022 (Maldonado et al., 2012). The sponge specimens were collected from depths of 3 to 5 meters using snorkeling and SCUBA diving techniques (Cleary et al., 2023). At each site, the sponges were sampled based on the purposive sampling criteria and subsequently rinsed with sterile ocean water to remove debris and epiphytic organisms (Williams et al., 2020).

Specimens were placed in labeled containers filled with sterile ocean water and transported to the Kenya Marine and Fisheries Research Institute (KMFRI) laboratories. Preservation methods were selected based on the intended further analyses: samples were frozen at -20°C for genetic studies and antimicrobial activity assessment, and preserved in 70% ethanol for morphological evaluations (Schöttner et al., 2013).

### Morphological characterization of the marine sponges

The sponge morphological identification relied on external morphology, encompassing the color, shape, and surface features (Hadi et al., 2016). The spicules and spongin fibers within the marine sponge skeleton were digested into small pieces of sponge tissue in bleach and observed under a Primo Star ZEISS image analyzer microscope with a coverslip. The microphotographs were taken using an Axio cam ERc5s digital camera (Carl Zeiss, Germany) (Jensen et al., 2011). The marine sponge specimens voucher numbers: BLSi 007, BRMu 004, and BLD 014 used in this study are preserved at the Kenya Marine and Fisheries Research Institute Museum. The marine sponge species was determined using the World Register of Marine Species (WoRMS) database (Costello et al., 2013). To identify the genus and species of marine sponges, the Sponge Identification Reference Book and the Porifera database list (http://www.marinespecies.org/porifera/) were employed (Sandes et al., 2024).

### DNA Extraction and PCR Amplification of the Mitochondrial Gene Cytochrome c Oxidase Subunit I (COI)

The salting out method was used for DNA extraction for the sponge specimens (Daniels et al., 2023). First, 200 µl of the extraction buffer and 5 µl of proteinase K (Thermo Fisher Scientific, Unites States) were added to the tissue. The mixture was incubated at 37°C overnight. Following the incubation, the proteinase K was added, and the mixture was spun down, and 450 µl of 3M sodium chloride (NaCl) was added and vortexed (Goldschmidt et al., 2014). The sample was then centrifuged at a speed of ≥10,000 g for 15 minutes to collect cell debris. The supernatant, containing DNA (1 ml), was transferred to a sterile 1.5 ml tube and mixed with frozen (-20°C) absolute ethanol to fill the tube (Marín et al., 2021). The DNA was allowed to precipitate overnight in a freezer at -20°C, followed by centrifugation at a speed of ≥10,000 g for 20 minutes. The pellet was washed thrice with 700 µl of 70% ethanol and then centrifuged at a speed of ≥10,000 g for another 20 minutes. The ethanol was pipetted out without disturbing the pellet, which was allowed to dry at room temperature (Holub et al., 2022). The DNA was then resuspended in 50 µl of deionized water (molecular grade) (Carl Roth, Germany) and its concentration and purity were assessed using a NanoDrop spectrophotometer (Thermo Fisher Scientific, United States).

The degenerate universal barcoding primers dgLCO1490 (GGT CAA CAA ATC ATA AAG AYA TYG G) and dgHCO2198 (TAA ACT TCA GGG TGA CCA AAR AAY CA), targeting the mitochondrial gene cytochrome c oxidase subunit I (COI) were used for amplification (Patantis et al., 2014). The PCR reaction mixture of 25 µL was prepared by mixing 12.5 µL of 2X PCR Master Mix (DreamTaq Green PCR, Thermo Fisher Scientific, United States), which included Taq polymerase, dNTPs, and buffer. Additionally, 1.0 µL of both the forward and reverse primers (10 µM) and 2.0 µL of template DNA (20 ng/µL) were added. The reaction volume was then brought to 25.0 µL by adding 8.5 µL of nuclease-free water. For PCR product verification, a 1.5% agarose gel (Sigma-Aldrich, Merck Millipore) was prepared in 1X TAE buffer, and stain G (SERVA Electrophoresis, Germany) was added (Timmers et al., 2022). A 5 µL aliquot of the PCR product was mixed with 1 µL of SYBR Green dye (Thermo Fisher Scientific, United States) and loaded into the gel wells. Distilled water was used as the negative control, while the DNA ladder (CSL-MDNA, Cleaver Scientific) was used as a DNA marker. The gel was run at 100V for 30 minutes, and the PCR products were visualized under UV light using a gel documentation system (ATTO, Tokyo, Japan).

The PCR products were treated with an ExoSAP treatment (Kanakaraj et al., 2011). The purified products were then shipped to Inqaba Biotec, a commercial sequencing service provider in South Africa, for Sanger sequencing.

### DNA Sequencing and Phylogenetic Analysis

Sequencing was conducted using the same primers (dgLCO1490 and dgHCO2198) that were used for PCR amplification. The reaction mixture consisted of the purified DNA template, sequencing primers, BigDye Terminator v3.1 Cycle Sequencing Kit, and buffer (SenGupta and Cookson, 2010). The prepared reaction mixture was subjected to cycle sequencing and then loaded onto an automated DNA sequencer (Applied Biosystems 3500XL Genetic Analyzer). The fluorescence data were interpreted using the FinchTV analysis software to generate the DNA sequence (Shuhaib and Hashim, 2023).

The raw sequence data from Sanger sequencing were processed using BioEdit software version 7.2. Low-quality bases were identified and removed to maintain data integrity. The sequences were compared against the nucleotide sequence database (GenBank) at the National Center for Biotechnology Information (NCBI) and Barcode of Life Database (BOLD) using the BLASTn algorithm to determine species identity. Additionally, the BOLD platform was utilized for sponge DNA barcoding, focusing on species-level identification through a curated reference library of standardized mitochondrial COI (cytochrome oxidase I) genetic markers.

Phylogenetic analysis was conducted using the Unipro UGENE software platform (https://ugene.net/). DNA sequences obtained from GenBank and BOLD databases were together with the newly obtained sequences, were imported in FASTA format and aligned using the MAFFT (Multiple Sequence Alignment using Fast Fourier Transform) algorithm (Zhang et al., 2018). The Maximum Likelihood method was employed to construct the phylogenetic tree, incorporating bootstrap resampling with 1,000 replications. The resulting phylogenetic tree was visualized using FigTree v1.4.4, with posterior probabilities displayed at the nodes to indicate the support for the branches (Zou et al., 2024).

### Crude Extracts Preparation for Antimicrobial Assays

The sponge samples were freeze-dried and ground into a fine powder using a mechanical grinder (Erngren et al., 2021). The powdered sponge material was subjected to solvent extraction using methanol, ethyl acetate, and dichloromethane to obtain bioactive compounds with varying polarities (Longo et al., 2025). The bioactive compounds recovery involved macerating the sponge powder in the respective solvents for 48 hours at room temperature with occasional agitation (Varijakzhan et al., 2021). The sponge extracts were then filtered using Whatman No. 1 filter paper (Sigma-Aldrich Company, St. Louis, Germany) and concentrated under reduced pressure using a rotary vacuum evaporator (BIOBASE Company, Jiangsu, China) at temperatures below 40°C to prevent thermal degradation of bioactive compounds (Mohd et al., 2023). The concentrated crude extracts were weighed and stored in sterile vials at 4°C until further analysis. The sponge extracts were assessed for their antimicrobial efficacy against test microorganisms: *Candida albicans* ATCC 10231, *Pseudomonas aeruginosa* ATCC 27853, *Escherichia coli* ATCC 25922, and *Staphylococcus aureus* ATCC 25923. All microbial strains utilized in this study were procured from the Kenya Medical Research Institute (KEMRI), Kenya.

### Assessment of *In Vitro* Antimicrobial Activity of Sponges’ Crude Extracts

The antimicrobial screening of the sponge crude extracts against the test microorganisms was conducted using the Kirby-Bauer disk diffusion method (Latifah et al., 2021a). Sterile Mueller-Hinton agar (MHA) (HiMedia, Mumbai, India) was utilized as the medium for bacterial susceptibility testing (Salam et al., 2023). The MHA was prepared following the manufacturer’s specifications (38.0 g in 1000 mL of distilled water). For turbidity standardization, the bacterial test organisms were cultured in Mueller-Hinton broth (MHB) at 37°C for 18 hours with continuous agitation at 150 revolutions per minute (rpm) to ensure uniform bacterial suspension (Jalal et al., 2023). The MHB (HiMedia, Mumbai, India) was prepared according to the manufacturer’s guidelines (21 g in 1000 mL of purified water).

For *C. albicans*, Potato Dextrose Agar (PDA) (HiMedia, Mumbai, India) was used. *C. albicans* broth cultures were incubated at 30°C for 48 hours in Potato Dextrose Broth (PDB), prepared as per the manufacturer’s protocol (24 g in 1000 mL of distilled water) for turbidity standardization. Microbial cultures were adjusted to a 0.5 McFarland turbidity standard, representing approximately 1.5 × 10^8^ CFU/mL for bacterial cells and 1 × 10^6^ CFU/mL for fungal cells, using sterile normal saline as the diluent. The normal saline was prepared by dissolving 9 g of NaCl in 1000 mL of deionized water.

MHA and PDA plates were inoculated with microbial test strains employing a sterile swab technique. Sponge crude extracts were solubilized in dimethyl sulfoxide (DMSO) to achieve working concentrations of 10 µL of 10 mg/mL sponge extracts in DMSO. Sterile paper discs (Oxoid, Thermo Scientific, United Kingdom) measuring 6 mm in diameter, were impregnated with 20 µL of sponge extracts and positioned on the inoculated agar plates (Manuel et al., 2021). Streptomycin (200 µg/ml) served as the positive control for bacterial strains, whereas fluconazole (10 mcg) was employed as the positive control for *C. albicans*, with DMSO functioning as the negative control (Domingos et al., 2024). Following incubation, plates were maintained at 37°C for bacterial strains and at 30°C for fungal strains, with measurements of inhibition zone diameters (IZD) conducted in millimeters using a digital caliper to evaluate antimicrobial efficacy (Gajic et al., 2022).

### Determination of the minimum inhibitory concentration (MIC)

The broth microdilution method was employed to evaluate the antimicrobial efficacy of the marine sponge extracts against the selected pathogenic microbial strains (Anteneh et al., 2021). This method involved the preparation of a two-fold serial dilution of the marine sponge extracts in nutrient broth for antibacterial screening and in PDB for antifungal assessment (Balouiri et al., 2016). The test tubes were systematically labelled (Tube 2, Tube 3, Tube 4, Tube 5, Positive Control (+ve), and Negative Control (-ve). A stock solution of the sponge extract (Tube A) was initially prepared for subsequent dilutions. A volume of 2 mL of sterile broth was dispensed into each labelled test tube, beginning with the negative control. For the serial dilution, 2 mL of the sponge extract from Tube A was transferred into Tube 2 and mixed thoroughly. Subsequently, the process was repeated sequentially for Tubes 3 to 5. The concentration gradient of the stock solution in the nutrient broth resulted in final concentrations of 0.625 mg mL^-1^, 1.25 mg mL^-1^, 2.5 mg mL^-1^, 5 mg mL^-1^, and 10 mg mL-1 of the sponge extracts (Júnior et al., 2021).

Following the preparation of the serial dilutions, 0.3 mL of the microbial suspension (*E. coli*, *S. aureus*, *P. aeruginosa*, and *C. albicans*), was introduced into all tubes except the negative control (Parvekar et al., 2020). The test tubes were then incubated under aerobic conditions at 37°C for 18 hours for bacterial strains and at 30°C for 48 hours for *C. albicans* to allow for microbial growth (Luo et al., 2021). After the incubation period, bacterial growth was assessed by measuring the optical density (OD) at 600 nm using a spectrophotometer (Shimadzu, Japan) (Mytilinaios et al., 2012). The minimum inhibitory concentration (MIC) was determined as the lowest concentration of the sponge extract that visibly inhibited bacterial growth (Kowalska and Dudek, 2021).

### Evaluation of the minimum bactericidal concentration (MBC) and minimum fungicidal concentration (MFC)

The minimum bactericidal concentration (MBC) and minimum fungicidal concentration (MFC) were determined through the subculturing of wells from the MIC assays onto fresh agar plates (Abedini et al., 2020). Specifically, the bacterial cultures were incubated for 18 hours at 37 °C (Tuttle et al., 2021), while *C. albicans* cultures were incubated at 30 °C for 48 hours (Casagrande Pierantoni et al., 2021). Following incubation, the MHA plates were examined for the presence of surviving organisms. If the MBC of the sponge extracts was less than or equal to the MBC of the standard drug, the extract’s activity was classified as either bactericidal or fungicidal (Giugliano et al., 2023).

### Gas Chromatography-Mass Spectrometry (GC-MS) of Sponge Crude Extracts

The dried sponge extracts were reconstituted in analytical-grade n-hexane of high purity to attain a final concentration of 1 mg/mL (Rahman et al., 2024). Subsequently, the reconstituted samples were subjected to filtration through a 0.22 µm membrane filter to eliminate any particulate contaminants. A 1 µL aliquot of the prepared extract was injected into the GC-MS system for analytical assessment (Puah et al., 2019). The GC-MS system was equipped with an HP-5MS capillary column (5% phenyl methylpolysiloxane, 30 m × 0.25 mm i.d. × 0.25 µm film thickness) (Czarnowski et al., 2024). High-purity helium served as the carrier gas at a constant flow rate of 1 mL/min, and the analysis was performed in splitless mode to enhance sensitivity. The injector temperature was maintained at 250°C, while the oven temperature program commenced at an initial temperature of 50°C, held for 2 minutes, followed by a ramp rate of 10°C/min up to a final temperature of 300°C, where it was sustained for 5 minutes (Oliver, 2017).

The mass spectrometer was operated in Electron Ionization (EI) mode at an electron energy of 70 eV, with ion source temperature of 230°C and a quadrupole temperature of 150°C. The mass scan range was configured from 50 to 600 m/z, with a solvent delay period of 3 minutes. The mass spectra of the compounds detected in this study were systematically compared against established reference spectra derived from the National Institute of Standards and Technology (NIST) libraries (Valdez et al., 2021). An analysis of retention times and molecular ion peaks was conducted to authenticate the identities of the compounds. The relative abundance of bioactive compounds was quantified through peak area integration, enabling a precise determination of their prevalence. Identified compounds were categorized according to their chemical classes, including alkaloids, terpenoids, steroids, and fatty acids. The results were compared with previously reported marine-derived bioactive compounds to assess their novelty and potential pharmaceutical applications (Mehbub et al., 2024).

## Data analysis

A two-way analysis of variance (ANOVA) was performed to determine the presence of statistically significant differences in the diameters of inhibition zones across various marine sponge extracts. The data were recorded in triplicate and presented as mean ± standard deviation (SD) for the inhibition zones. Statistical tests of means sharing identical superscripts (*) within the column were considered not significantly different from one another, as determined through Fisher’s Least Significant Difference (LSD) test at a 95% confidence level, following a post hoc analysis conducted using Minitab software version 21.4.1. Differences were classified as statistically significant at P < 0.05 (α = 0.05).

The mass spectra of each compound analyzed by GC-MS were compared with the National Institute of Standards and Technology (NIST) spectral library for identification.

## RESULTS

### Morphology and taxonomy of marine sponges

These sponge samples were collected from Sii Island, Mundini, and Ras Kiromo study sites and taxonomically identified as *Biemna fistulosa, Callyspongia diffusa,* and *Haliclona fascigera*, respectively.

Upon removal from their natural habitat, the sponges exhibited significant morphological changes, including pronounced shrinkage and increased fragility. Additionally, a notable decline in their vibrant coloration was observed (Figure 1).

**Figure 1:**
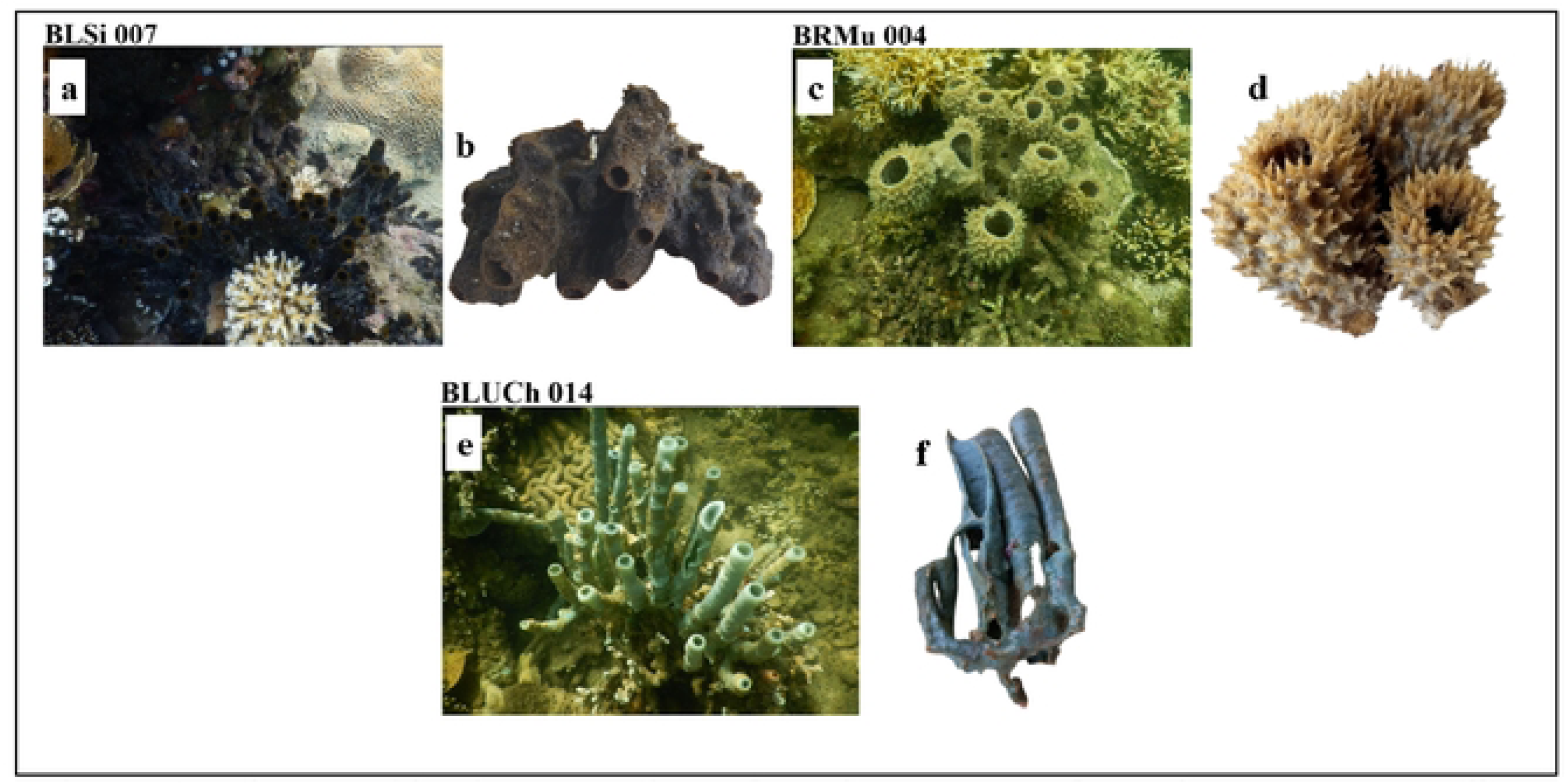
Photographic documentation of marine sponges from the Kenyan coastline, presented as follows: **(a)** *Biemna fistulosa* (Voucher specimen (BLSi 007) (in-situ) **(b)** *B. fistulosa* (detached) **(c)** *Callyspongia diffusa* (Voucher specimen BRMu 004) (in-situ) **(d)** *C. diffusa* (detached) **(e)** *Haliclona fascigera* (Voucher specimen BLUCh 014) (in-situ) **(f)** *Haliclona fascigera* (detached).

*Biemna fistulosa* sponge exhibited a robust, spongy and fibrous texture, with a growth pattern that encrusted and branched upwards (Figure 2). Initially, it presented a white-grey hue underwater, which upon exposure to air, turned black. The species, *B. ehrenbergi*, typically inhabited sandy shores and mangrove lagoons. Its structure was characterized by densely packed diactinal curved in styles (685.0-970.9-1235.7 x 7.9-19.0-32.8 μm). These spicules formed an irregular network, creating polygonal patterns. Additionally, microscleres such as C-shaped sigmas (30.4-41.0-50.3 μm), were also observed.

**Figure 2:**
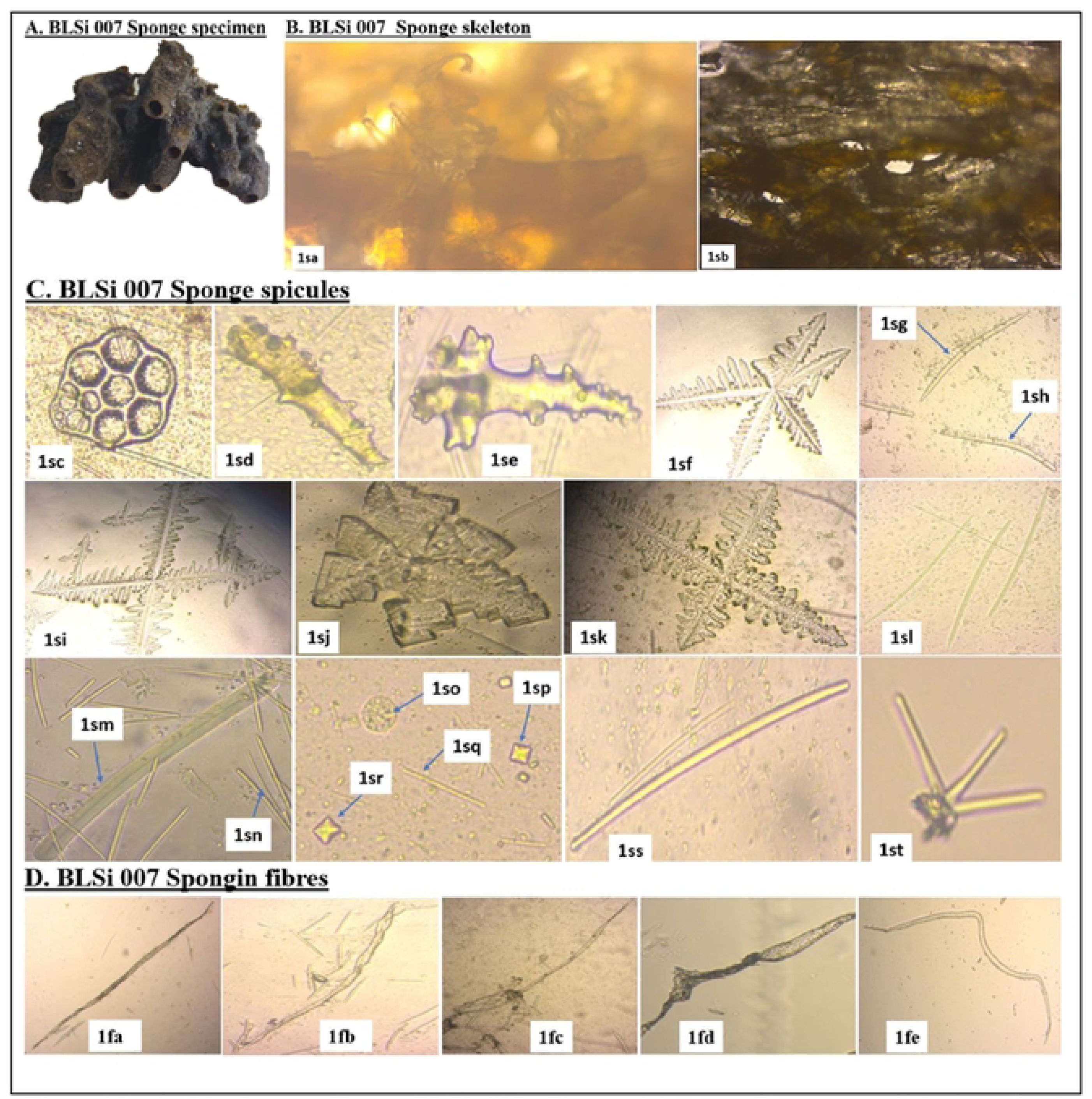
**A**. **Marine poriferan BLSi 007**, *Biemna fistulosa*; **B. sponge skeleton**: *(1sa):* perpendicular section; and *(1sb):* a tangential section (40x magnification); **C. sponge spicules:** *(1sc):* synapta plates; *(1sd)* and *(1se):* megascleres acanthostyles; *(1sf):* pentactines megascleres with digits at the tentacles; *(1sg):* curved oxeas; *(1sh):* curved styles; *(1si)* and *(1sk):* stauractines megascleres with digits at the tentacles; *(1sj):* pentactines megascleres with digits at the tentacles; *(1sl):* styles; *(1sm):* tabulated strongyles; *(1sn):* raphides; *(1so):* Microbiota (*Coscinodiscus radiatus*); *(1sp)* and *(1sr):* sterrasters; *(1sq):* microstrongyles; *(1ss):* strongyles; and *(1st):* dendroclones (unique to extinct sponge); **D. spongin fibers:** *(1fa):* simple elongated spongin fibers; *(1fb):* simple irregular spongin fibers; *(1fc):* Spongin fiber forming a fiber network on one end; *(1fd):* Spongin fiber with an irregular shape; *(1fe):* spongin fiber with a curved structure.

*Callyspongia diffusa* also called the diffuse rope sponge exhibited a main skeleton composed of an isodictyl reticulation of collagenous spongin fibers, with diactinal spicules forming triangular meshes in the skeleton (Figure 3). The *C. diffusa* megascleres were represented by anchorates, acanthostyles (209.0-224.5-267.0 x 5.7-7.9-11.9 μm) and strongyles (163.9-178.2-179.3 x 4.9-7.4-10.7 μm), while the microscleres took the form of sterrasters. Additionally, the sponge was characterized by numerous raphides (149.3-168.6-279.8 μm).

**Figure 3:**
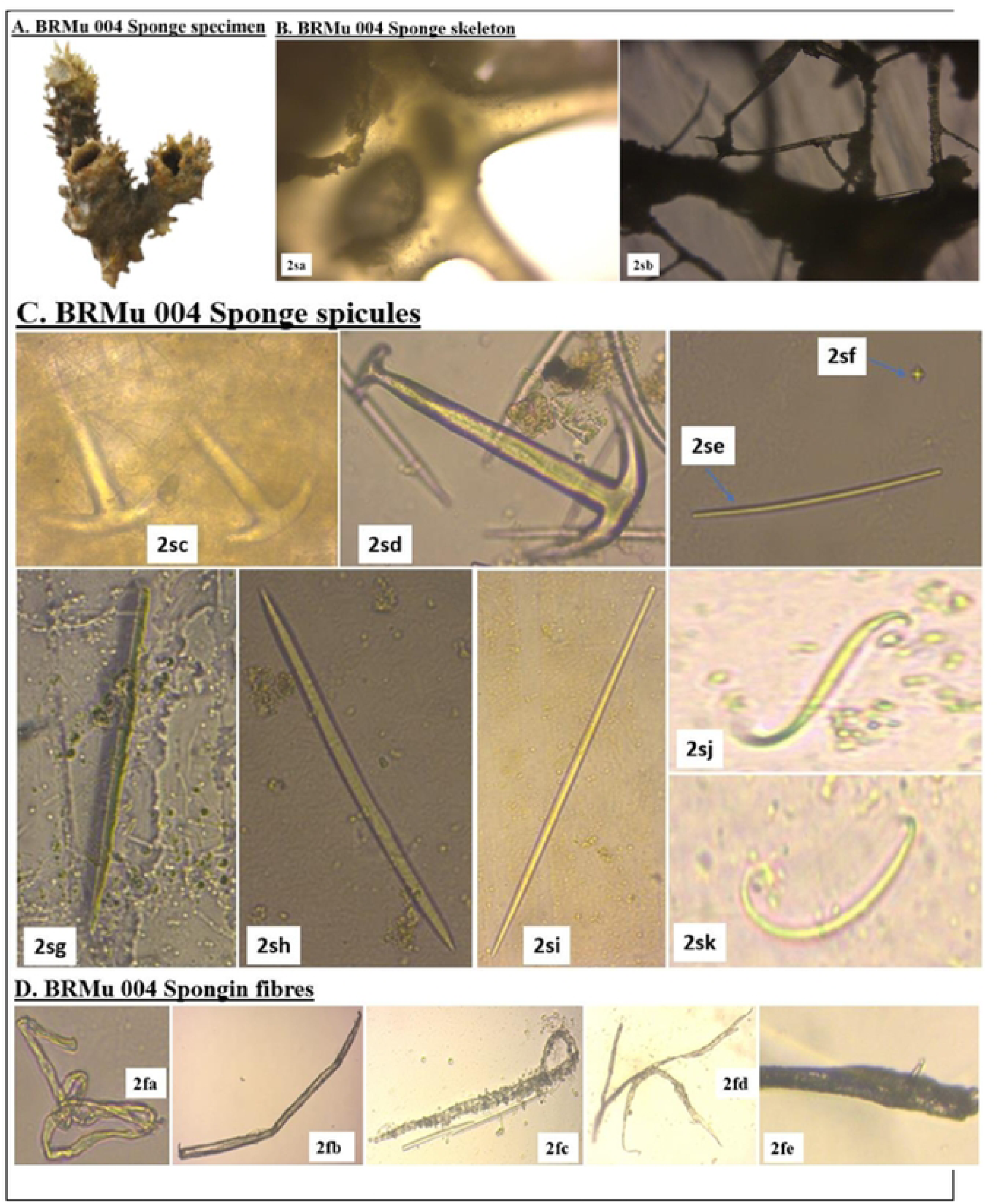
**A**. **Marine poriferan BRMu 004**, *Callyspongia diffusa*; **B. sponge skeleton**: *(2sa):* perpendicular section; and *(2sb):* a tangential section (40x magnification); **C. sponge spicules:** *(2sc)* and *(2sd):* anchorates; *(2se):* strongyles; *(2sf):* sterrasters; *(2sg):* acanthostyles; *(2sh):* curved oxeas; *(2si):* styles; *(2sj):* S sigmas; and *(2sk):* C sigmas; **D. spongin fibers:** *(2fa):* twisted thick spongin fibers; *(2fb):* Spongin fibers with bent thickened cell walls; *(2fc):* spongin fibers with a complete bent (*Microcoleus vaginatus* attaching on the surface); *(2fd):* spongin fibers with an anastomosing system; and *(2fe):* spongin fibers with hard collagen material (spicules protruding).

The sponge *Haliclona fascigera* possessed a choanosomal skeleton, characterized by an isodictyal network of spicules intertwined with nodal spongin fibers, creating polygonal patterns (Figure 4). Its primary structural megascleres were exclusively oxeas (428-497.9-634.1 x 8.3-13.1-20.0 μm) and were embedded within spongin fibers. A significant number of raphides (114.2-163.0-179.3 μm), microstrongyles (128.0-148.0-178.4 μm) and synapta plates were observed. *H. fascigera* typically inhabited the shallow, sandy bottoms of lagoons and the vicinities of coral reefs.

**Figure 4:**
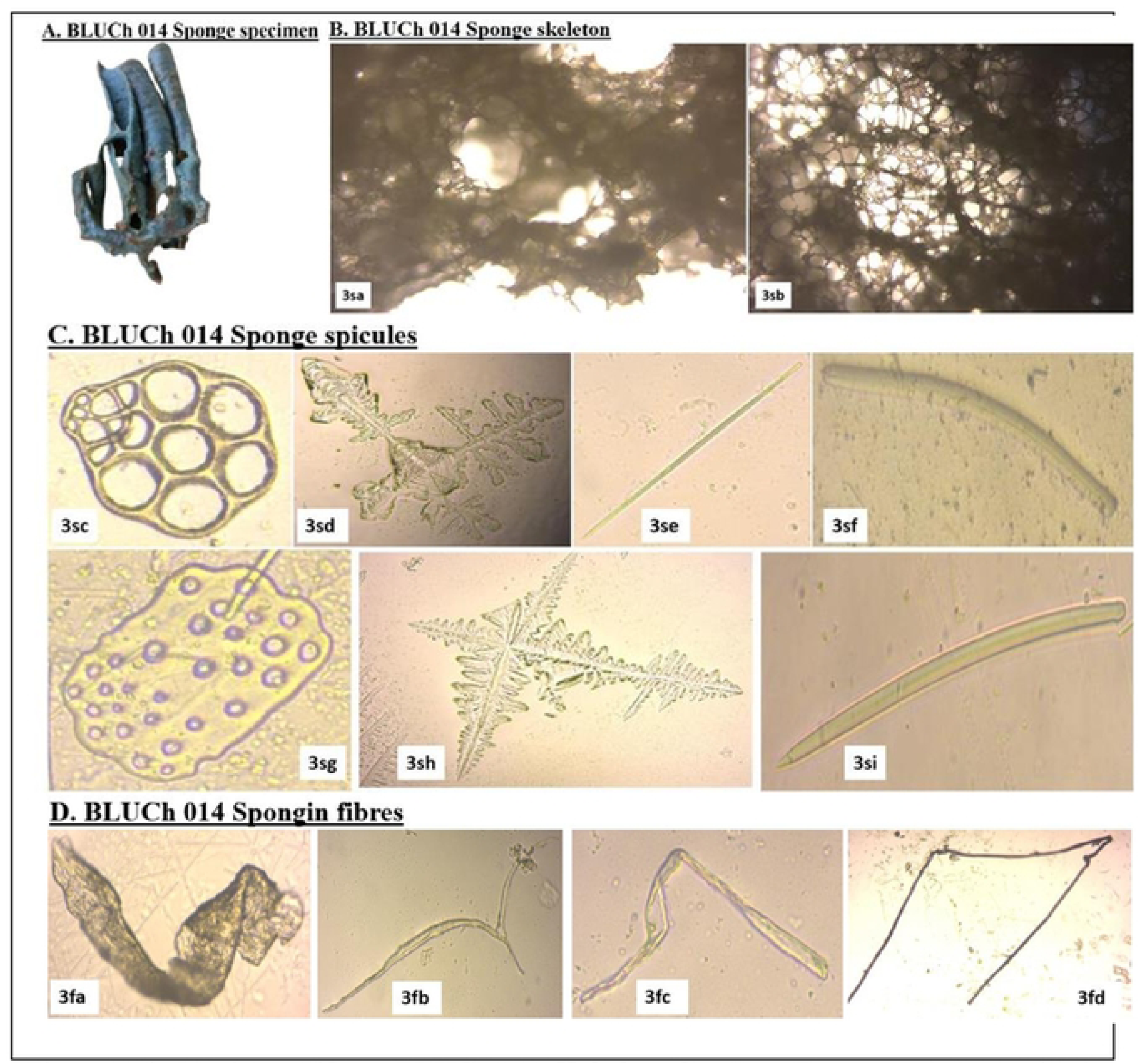
**A. Marine poriferan BLUCh 014**, *Haliclona fascigera*; **B. sponge skeleton**: *(3sa):* perpendicular section; and *(3sb):* a tangential section (40x magnification); **C. sponge spicules:** *(3sc):* synapta plates; *(3sd)* and *(3sh):* stauractines megascleres with digits at the tentacles; *(3se);* styles; *(3sf):* tuberculated curved strongyles; *(3sg):* large plates of calcareous deposits; *(3si):* styles; **D. spongin fibers:** *(3fa):* spongin fibers with a thick flat structure; *(3fb):* branched spongin fibers; *(3fc):* twisted spongin fibers with an open transparent lumen; *(3fd):* spongin fibers with thickened cell walls and a smooth transparent lumen

### Taxonomic Affiliation of the Marine Sponges

The genomic DNA extracted from marine sponge samples had NanoDrop quantifications ranging between 84.9 and 245.3 µg/mL. PCR amplification of the cytochrome c oxidase subunit I (COI) gene yielded amplicons within the expected size range of approximately 650 to 700 base pairs (bp). All sponge specimens were classified within the phylum Porifera and identified as belonging to the genera *Callyspongia, Haliclona*, and *Biemna* based on percentage sequence similarities. A comparative analysis of the newly obtained mitochondrial Cytochrome Oxidase Subunit 1 (CO1) sequences (PQ329108, PQ997929, and PQ997931), conducted using BLASTn search against the GenBank database, showed sequence similarities of ≥99% with existing entries in the nucleotide sequence repository (Table 1). Each sponge sample formed a distinct sub-cluster (representing a specific genus), with a bootstrap support value of 100% (Table 1 and Figure 5). The marine sponge specimens BRMu 004 (PQ329108) from Mundini and BLUCh 014 (PQ997929) from Ras Kiromo were closely related to *Callyspongia diffusa* and *Haliclona fascigera*, respectively. These specimens had sequence identities of 100% and 99%, respectively (Table 1), and formed distinct subclusters, each supported by a bootstrap value of 100%, as depicted in the phylogenetic tree (Figure 5). The sponge specimen BLSi 007 (PQ997931) from Sii Island was closely related to the known sponge species *Biemna fistulosa*, with a sequence identity of 98%, and formed a subcluster with a bootstrap support value of 100% (Table 1 and Figure 5).

**Figure 5:**
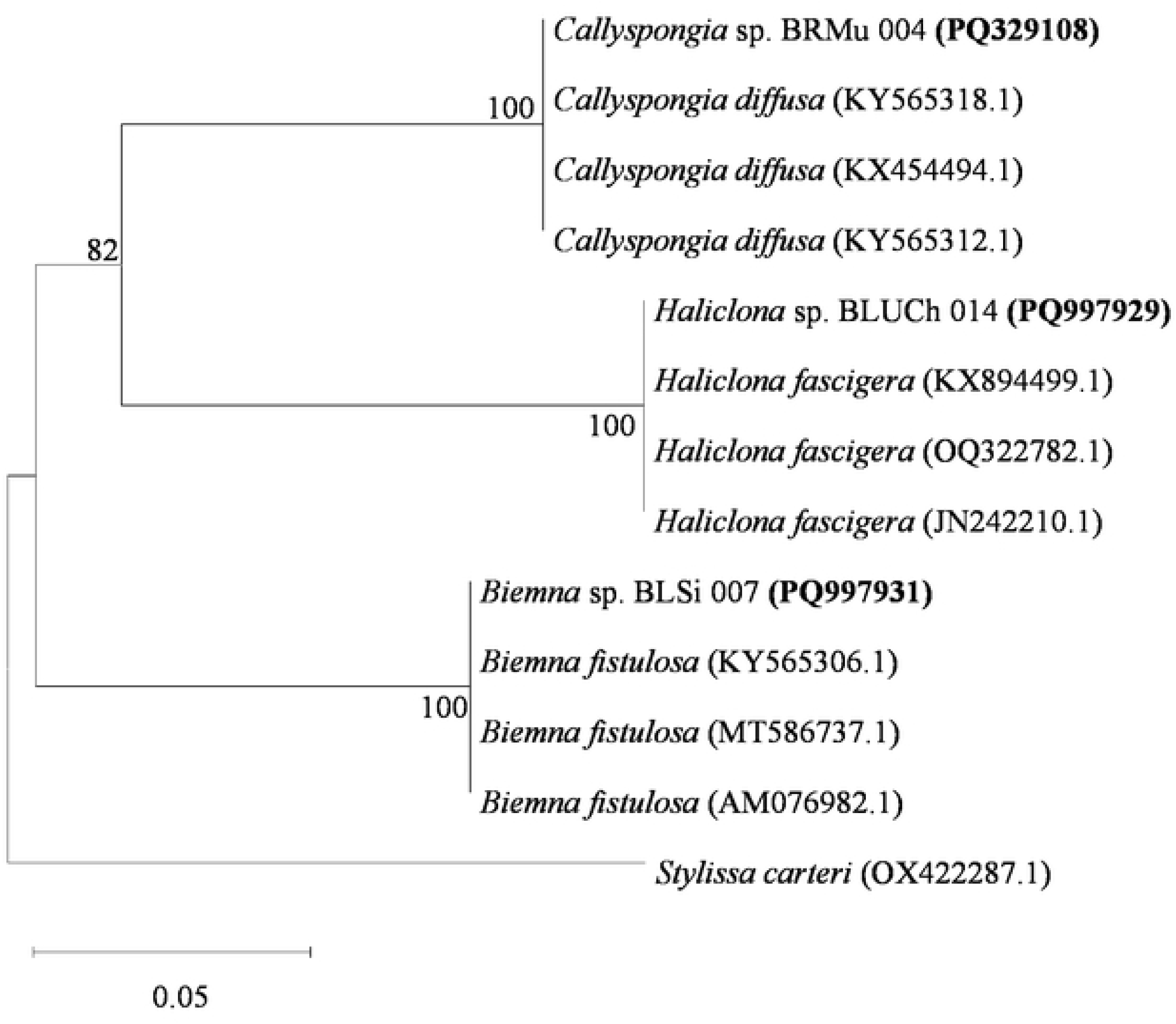
Phylogenetic Relationships of Metazoan CO1 Sequences with Closely Related Sponge Species. The phylogenetic tree was rooted using *Stylissa carteri* (OX422287.1). Bootstrap values exceeding 50%, derived from 1000 replications, are indicated at the branch nodes. The scale bar represents 0.05 substitutions per nucleotide.

**Table 1:** Taxonomic affiliation of marine metazoan sponges with their closest phylogenetic relatives.

| Sample ID | Accession No. | Location | Closest taxonomic affiliation | Isolation Source | Country | % ID |
| --- | --- | --- | --- | --- | --- | --- |
| BLSi 007 | PQ997931 | Sii Island (4°40'46.1"S, 39°17'01.9"E) | *Biemna fistulosa* (AM076982.1) | Coral reefs (Núñez Pons et al., 2017) | United States | 98 |
| BRMu 004 | PQ329108 | Mundini (4°39'10.0"S, 39°21'41.0"E) | *Callyspongia diffusa* (KX454494.1) | Indo-Pacific region (Pallela & Janapala, 2013) | India | 100 |
| BLUCh 014 | PQ997929 | Ras Kiromo (4°38'45.5"S, 39°19'23.4"E) | *Haliclona fascigera* (OQ322782.1) | Island (Handayani et al., 2015) | Indonesia | 99 |

### *In Vitro* antibiotic and antifungal activity of the marine sponge crude extracts

The marine sponge extracts were evaluated for their effectiveness against human pathogenic strains (*Candida albicans,* ATCC 10231, gram-negative bacteria *Pseudomonas aeruginosa* ATCC 25923 and *Escherichia coli* ATCC 25922 and the gram-positive bacterium *Staphylococcus aureus* ATCC 27853 (Tables 2, 3, and 4)). All the three crude organic extracts from the marine sponges (*B. fistulosa, C. diffusa* and *H. fascigera*) showed significant antimicrobial activity against at least one of the four tested microorganisms, exceeding the efficacy of the positive control (P < 0.05) (Tables 2, 3 and 4). The methanolic extracts from the sponges *B. fistulosa* (28.00±3.5 mm) and *H. fascigera* (28.33±3.8 mm) exhibited a broad spectrum of antibacterial activity against *E. coli* that was higher than the positive control (27.67±0.9 mm) (Table 3). The inhibitory activity of ethyl acetate extracts *C. diffusa)* against *P. aeruginosa* was observed to be higher (29.33±2.4 mm) compared to that of the positive control (26.67±0.7 mm) (Table 4).

**Table 2:**
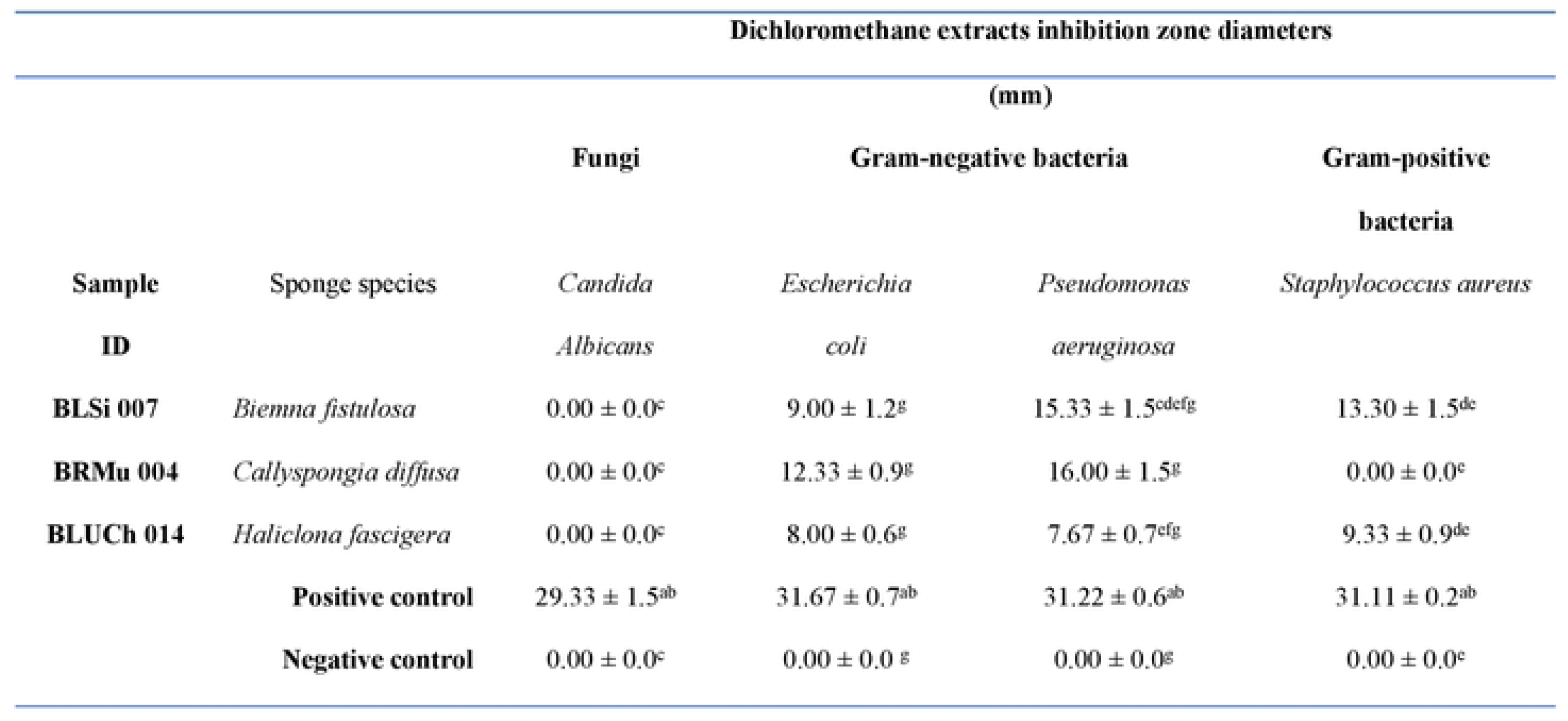
Antimicrobial activities of dichloromethane organic crude extracts from the selected marine sponges against the tested human pathogens.

**Table 3:** Antimicrobial activities of methanolic organic crude extracts from the selected marine sponges against the tested human pathogens.

| Methanolic extracts inhibition zone diameters |  |  |  |  |  |
| --- | --- | --- | --- | --- | --- |
|  |  | (mm) |  |  |  |
| Sample ID | Sponge species | Fungi | Gram-negative bacteria |  | Gram-positive bacteria |
|  |  | *Candida Albicans* | *Escherichia coli* | *Pseudomonas aeruginosa* | *Staphylococcus aureus* |
| BLSi 007 | *Biemna fistulosa* | 15.67 ± 1.2<sup>abc</sup> | 28.00 ± 3.5<sup>abcd</sup> | 19.67 ± 1.2<sup>abc</sup> | 25.33 ± 0.9<sup>abc</sup> |
| BRMu 004 | *Callyspongia diffusa* | 9.67 ± 2.2<sup>cd</sup> | 0.00 ± 0.0<sup>d</sup> | 0.00 ± 0.0<sup>c</sup> | 0.00 ± 0.0<sup>d</sup> |
| BLUCh 014 | *Haliclona fascigera* | 10.33 ± 0.9<sup>abc</sup> | 28.33 ± 3.8<sup>abcd</sup> | 15.67 ± 0.7<sup>ab</sup> | 24.67 ± 1.2<sup>abc</sup> |
|  | Positive control | 25.67 ± 1.2<sup>ab</sup> | 27.67 ± 0.9<sup>a</sup> | 26.67 ± 0.7<sup>ab</sup> | 31.11 ± 0.2<sup>abc</sup> |
|  | Negative control | 0.00 ± 0.0<sup>d</sup> | 0.00 ± 0.0<sup>d</sup> | 0.00 ± 0.0<sup>c</sup> | 0.00 ± 0.0<sup>d</sup> |

**Table 4:**
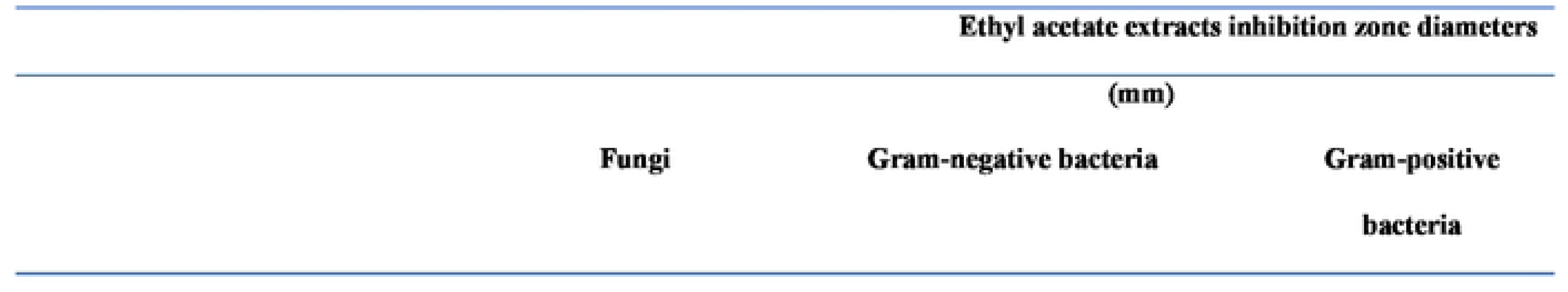
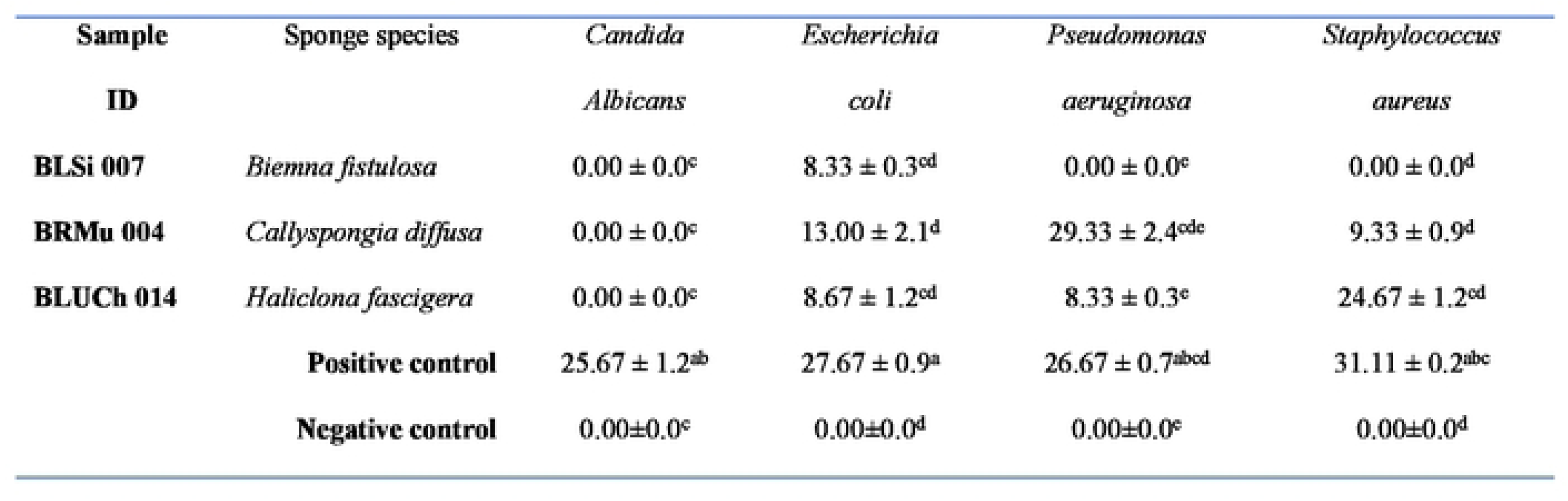
Antimicrobial activities of ethyl acetate organic crude extracts from the selected marine sponges against the tested human pathogens.

Methanolic extracts of *B. fistulosa* exhibited a higher (15.67±1.2mm) antifungal activity against *C. albicans* compared to the methanolic extract of *H. fascigera* (10.33±0.9 mm) and *C. diffusa* (9.67±2.2 mm) (Table 3). None of the dichloromethane and ethyl acetate extracts of the sponges exhibited antifungal activity against *C. albicans* (Table 2 and Table 4).

The marine sponges were coded based on their lifeform color and the study site of collection. The prefixes BL, BR, and BLU denoted black, brown, and blue sponges, respectively. These were followed by Si, Mu, and Ch, representing Sii Island, Mundini, and Ras Kiromo, respectively, along with the specimen number assigned to each sponge from the Kenyan coastline. Streptomycin was used as the positive control for bacterial strains, fluconazole as the positive control for *Candida albicans*, and Dimethyl Sulfoxide (DMSO) as the negative control. Data are expressed as means ± standard deviations (SD) from three replicates. Within each group, means sharing identical superscript letters indicate no significant difference at a 95% confidence level (α = 0.05), as determined by Fisher’s Least Significant Difference (LSD) test.

The tropical marine sponges were coded based on their lifeform color and collection site. The prefixes BL, BR, and BLU represented black, brown, and blue sponges, respectively, followed by Si, Mu, and Ch, indicating Sii Island, Mundini, and Ras Kiromo sampling sites. Each sponge was assigned a specimen number corresponding to its collection from the Kenyan coastline. Streptomycin served as the positive control for bacterial strains, fluconazole for *Candida albicans*, and Dimethyl Sulfoxide (DMSO) as the negative control. Data are presented as means ± standard deviations (SD) from three replicates. Within each group, means sharing the same superscript letter indicate no significant difference at a 95% confidence level (α = 0.05), as determined by Fisher’s Least Significant Difference (LSD) test.

The metazoan marine sponges were coded based on their lifeform color and collection site. The prefixes BL, BR, and BLU indicated black, brown, and blue sponges, respectively, followed by Si, Mu, and Ch, denoting Sii Island, Mundini, and Ras Kiromo sites. Each specimen was assigned a unique number corresponding to its collection from the Kenyan coastline. Streptomycin was employed as the positive control for bacterial strains, fluconazole for *Candida albicans*, and Dimethyl Sulfoxide (DMSO) as the negative control. Data are presented as means ± standard deviations (SD) from three replicates. Within each group, means with identical superscript letters signify no significant difference at a 95% confidence level (α = 0.05), as determined by Fisher’s Least Significant Difference (LSD) test.

### Evaluation of the sponge extracts for MIC, MBC and MFC

The Minimum Inhibitory Concentrations (MICs) of the sponge extracts were evaluated, demonstrating a range of values from 0.625 mg/ml to 10 mg/ml. Notably, streptomycin exhibited the lowest MIC values, with measurements of 1.94 mg/ml against *P. aeruginosa* and 1.36 mg/ml against *E. coli* (Table 5). The lowest MIC value (0.53±0.01 mg/ml) of methanolic extract of isolate *H. fascigera* was observed against *E. coli* compared to the standard drug/control (1.36±0.00 mg/ml) (Table 5). The MBC for all reference drugs was established at 2.5 mg/ml.

**Table 5:**
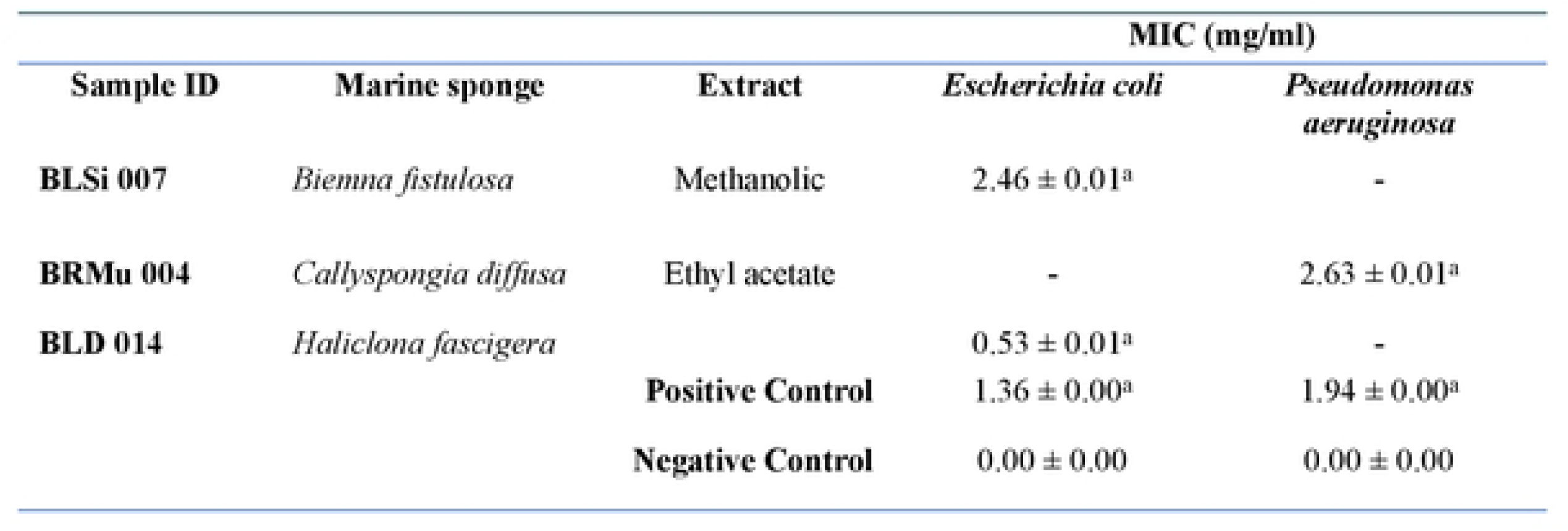
Minimum inhibitory concentrations (MIC) of dichloromethane, methanolic and ethyl acetate organic crude extracts of the selected marine sponges against the tested human pathogens.

The organic crude extract obtained from *H. fascigera* demonstrated the most potent bactericidal activity, with a minimum bactericidal concentration (MBC) of 1.25 mg/mL, exceeding the efficacy of the standard drug streptomycin as well as the extracts derived from *C. diffusa* and *B. fistulosa*. Specifically, the MBC values of the sponge extracts were recorded as 2.5 mg/mL for *C. diffusa* and 5 mg/mL for *B. fistulosa*. In comparison, the MBC of the streptomycin reference drug was established at 2.5 mg/mL concentration. The methanolic extract of *H. fascigera* demonstrated the most potent fungicidal activity, with a minimum fungicidal concentration (MFC) of 2.5 mg/mL. Notably, the extracts from the both sponges *C. diffusa* and *B. fistulosa* exhibited an MFC of 10 mg/mL. The fluconazole reference drug showed an MFC comparable to that of *H. fascigera*, with a value of 2.5 mg/mL concentration.

The tropical marine sponges were coded based on their lifeform color and collection site. The prefixes BL, BR, and BLU represented black, brown, and blue sponges, respectively, followed by Si, Mu, and Ch, denoting Sii Island, Mundini, and Ras Kiromo sampling sites. Each specimen was assigned a unique identification number corresponding to its collection location along the Kenyan coastline. Streptomycin was utilized as the positive control for bacterial strains, fluconazole served as the positive control for *Candida albicans*, and Dimethyl Sulfoxide (DMSO) functioned as the negative control. Data are presented as means ± standard deviations (SD) from three replicates. Within each group, means with identical superscript letters indicate no significant difference at a 95% confidence level (α = 0.05), as determined by Fisher’s Least Significant Difference (LSD) test.

### GC-MS spectral analysis of the crude extract of the marine sponge extracts

The GC-MS analysis of the extracts from *B. fistulosa, C. diffusa,* and *H. fascigera* generated a spectral profile and chemical structures of the detected compounds (Table 6). The peak numbers in the chromatograms correspond to the identified compounds (Figures 6–8 and Table 7). The GC-MS chromatogram data identified a total of 114 compounds across the three sponge extracts (BLSi 007, BRMu 004, BLUCh 014). These compounds belong to 39 distinct chemical classes (Table 6)

**Figure 6:**
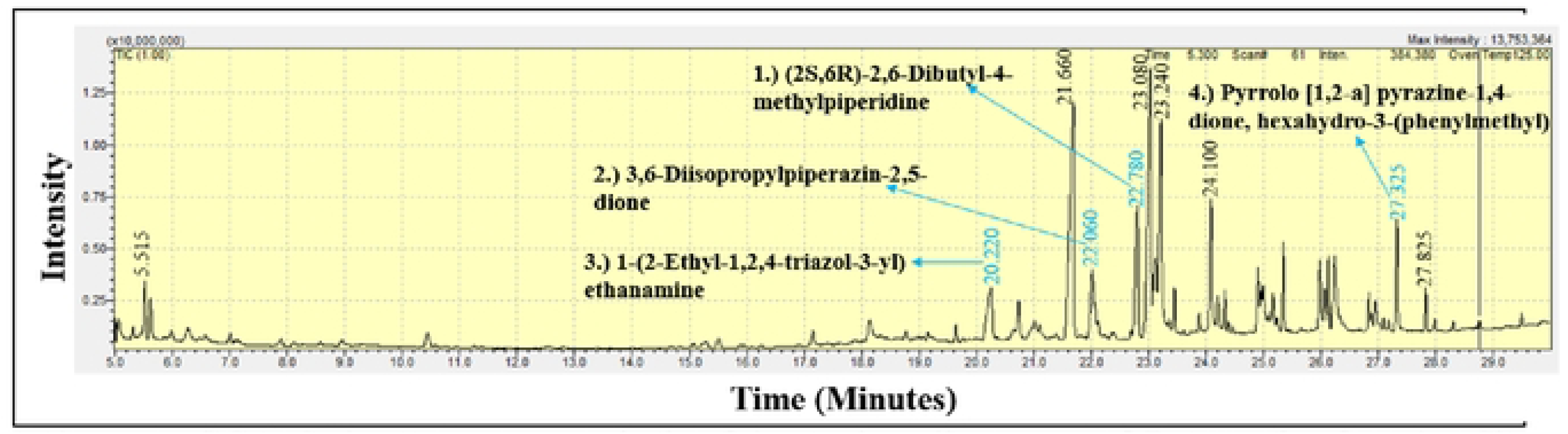
GC-MS chromatogram analysis of the methanolic extract of *Biemna fistulosa* (BLSi 007), highlighting four potent bioactive compounds

**Table 6:**
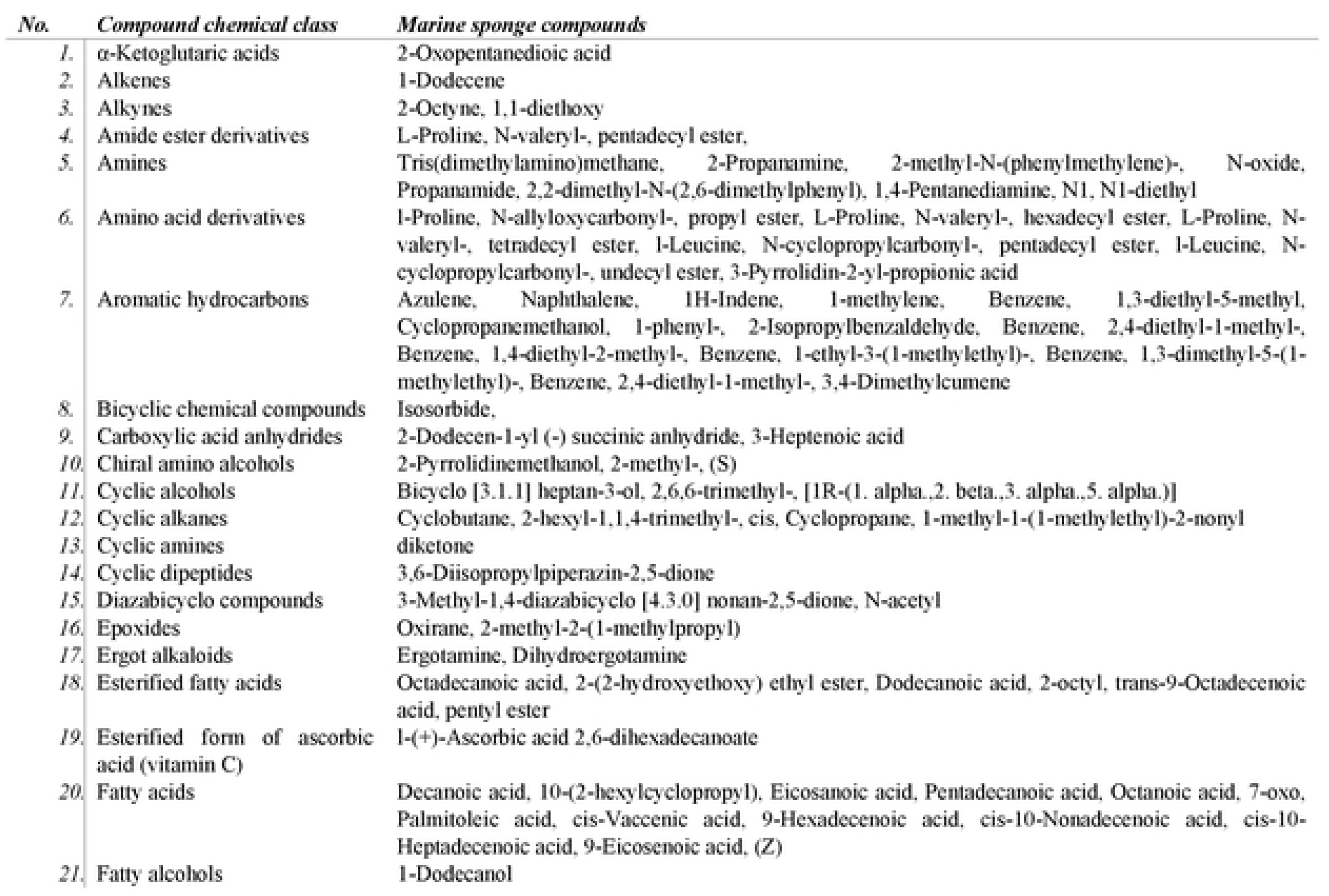
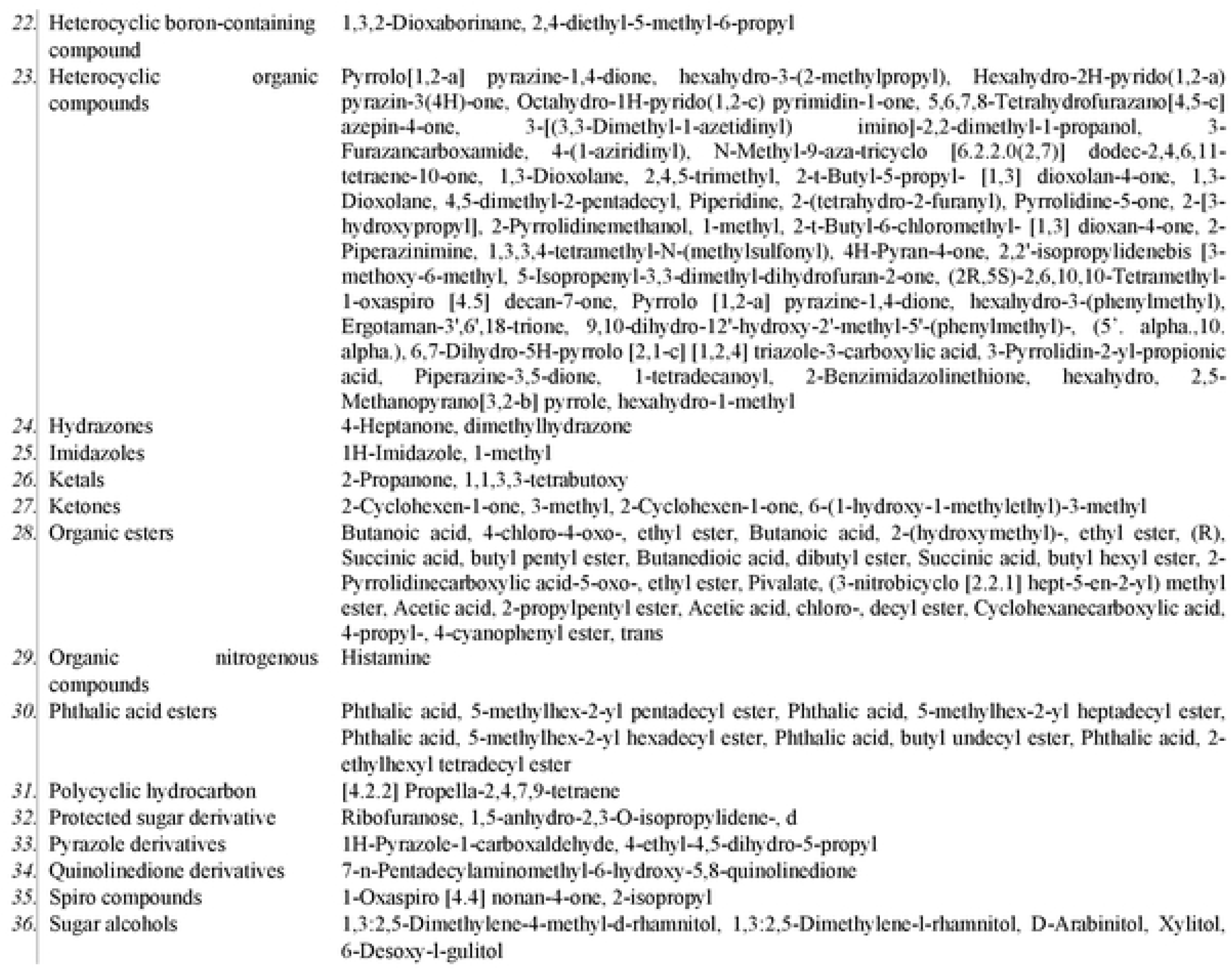
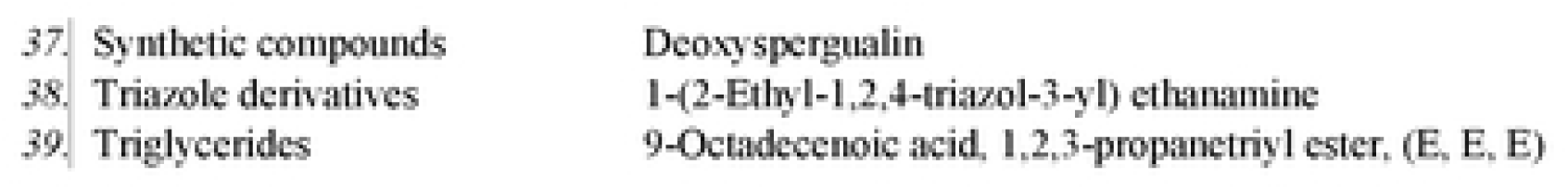
Classification of marine sponge compounds identified via GC-MS in organic extracts from *Biemna fistulosa, Callyspongia diffusa,* and *Haliclona fascigera* collected from Kenyan waters.

**Table 7:**
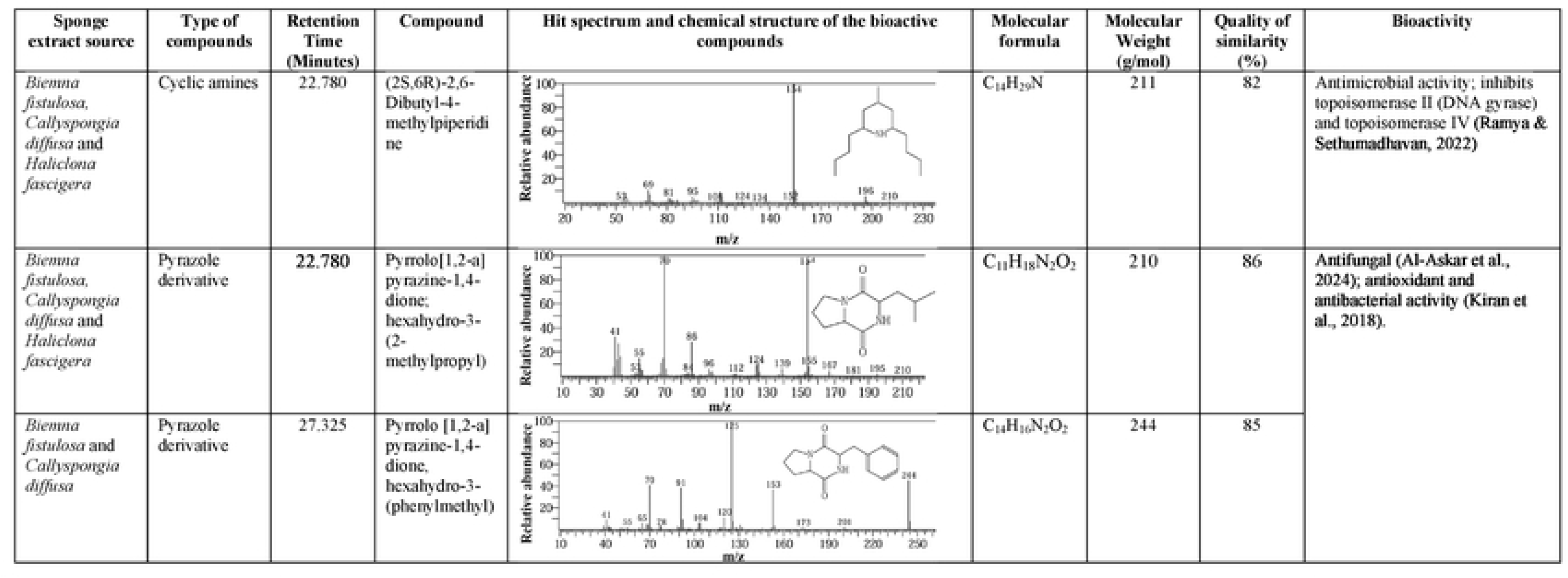
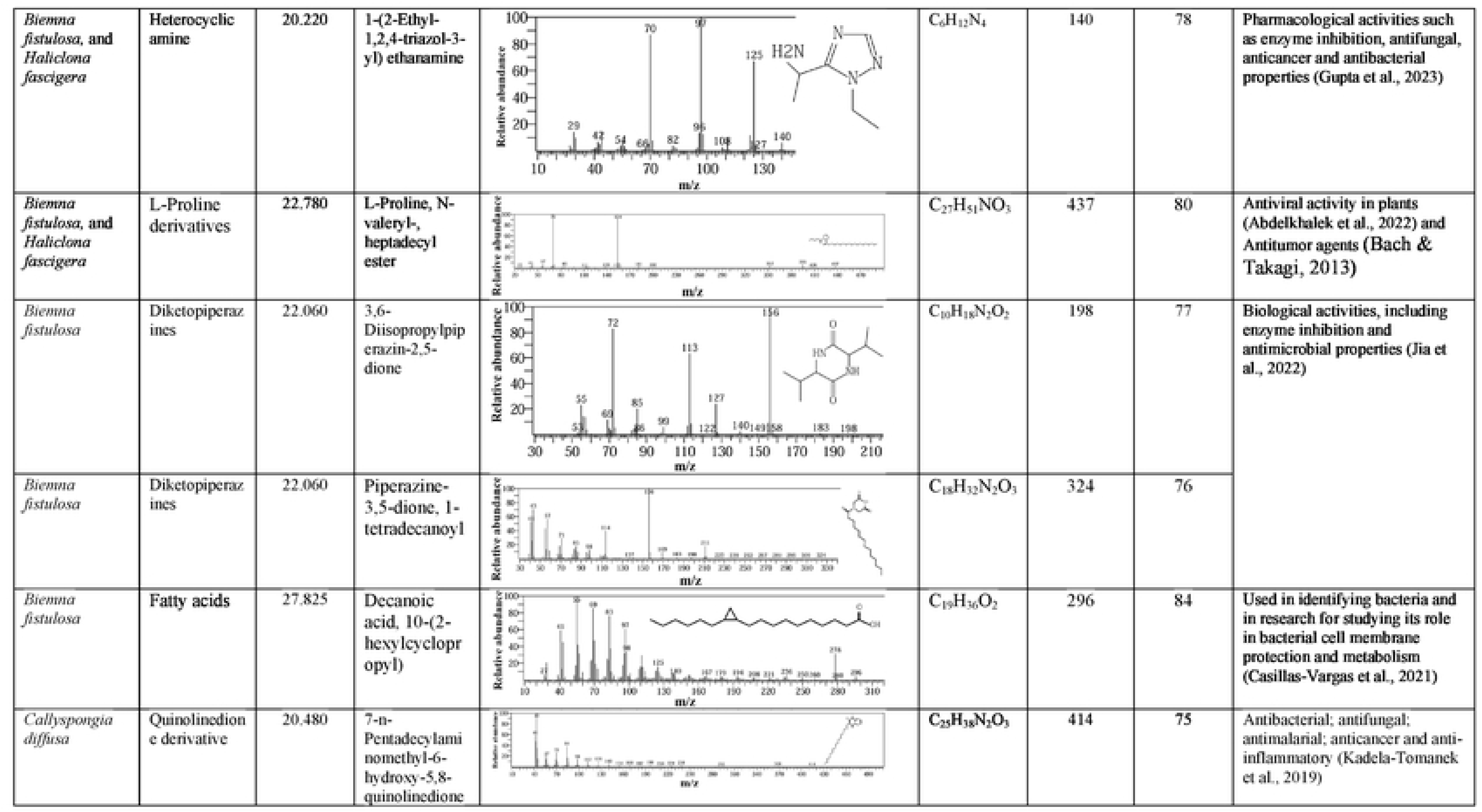
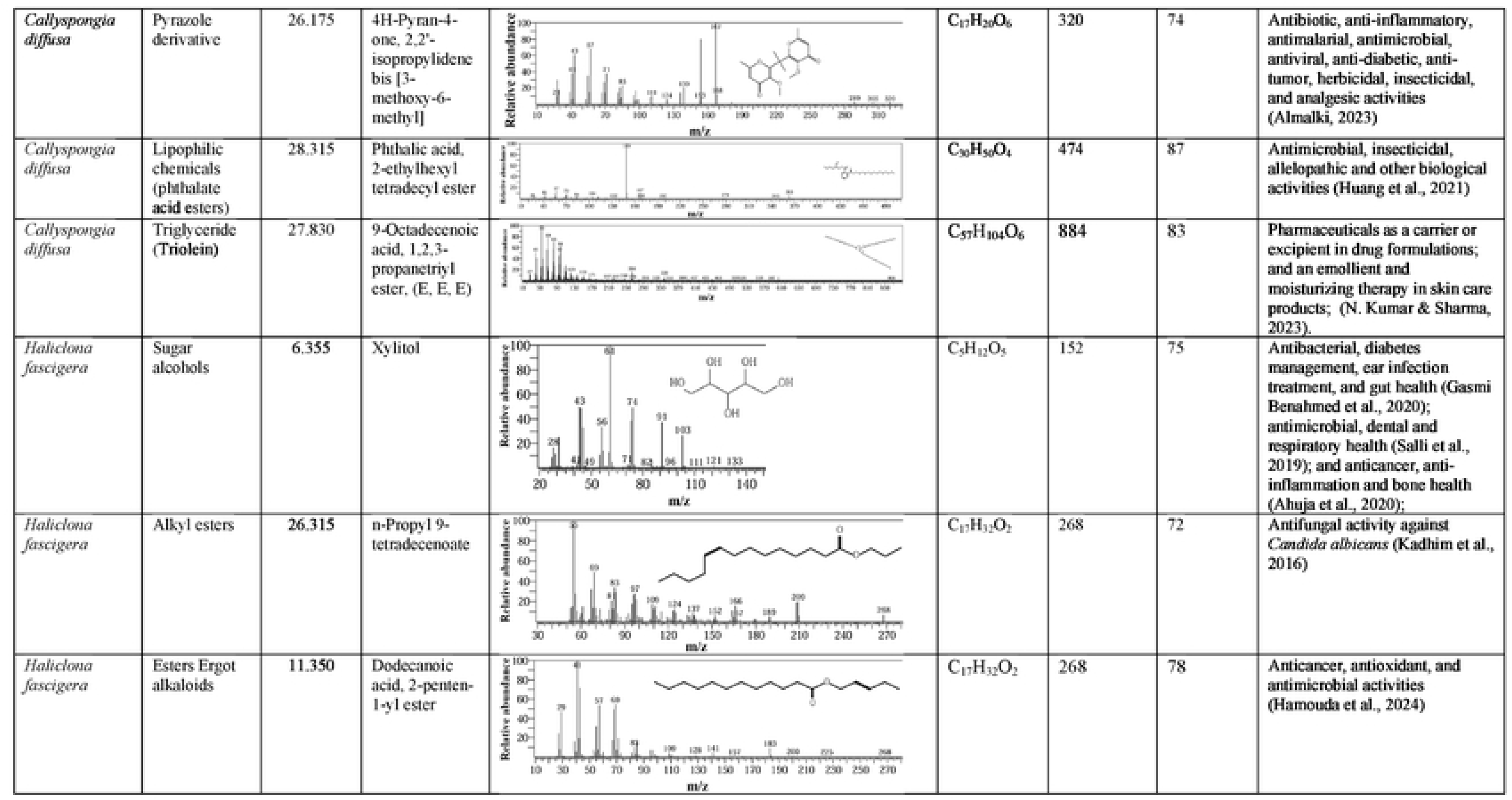
Characteristics and antimicrobial activity of selected sponges’ natural products identified in the GC-MS analysis of *Biemna fistulosa, Callyspongia diffusa,* and *Haliclona fascigera* organic extracts.

| Sponge extract source | Type of compounds | Retention Time (Minutes) | Compound | Hit spectrum and chemical structure of the bioactive compounds | Molecular formula | Molecular Weight (g/mol) | Quality of similarity (%) | Bioactivity |
| --- | --- | --- | --- | --- | --- | --- | --- | --- |
| *Biemna fistulosa*,<br>*Callyspongia diffusa* and<br>*Haliclona fascigera* | Cyclic amines | 22.780 | (2S,6R)-2,6-Dibutyl-4-methylpiperidine |  | C<sub>14</sub>H<sub>29</sub>N | 211 | 82 | Antimicrobial activity; inhibits topoisomerase II (DNA gyrase) and topoisomerase IV (Ramya & Sethumadhavan, 2022) |
| *Biemna fistulosa*,<br>*Callyspongia diffusa* and<br>*Haliclona fascigera* | Pyrazole derivative | 22.780 | Pyrrolo[1,2-a]pyrazine-1,4-dione; hexahydro-3-(2-methylpropyl) |  | C<sub>11</sub>H<sub>18</sub>N<sub>2</sub>O<sub>2</sub> | 210 | 86 | Antifungal (Al-Askar et al., 2024); antioxidant and antibacterial activity (Kiran et al., 2018). |
| *Biemna fistulosa* and<br>*Callyspongia diffusa* | Pyrazole derivative | 27.325 | Pyrrolo [1,2-a]pyrazine-1,4-dione, hexahydro-3-(phenylmethyl) |  | C<sub>14</sub>H<sub>16</sub>N<sub>2</sub>O<sub>2</sub> | 244 | 85 |  |
| *Biemna fistulosa*, and *Haliclona fascigera* | Heterocyclic amine | 20.220 | 1-(2-Ethyl-1,2,4-triazol-3-yl) ethanamine | | $C_6H_{12}N_4$ | 140 | 78 | Pharmacological activities such as enzyme inhibition, antifungal, anticancer and antibacterial properties (Gupta et al., 2023) |
| *Biemna fistulosa*, and *Haliclona fascigera* | L-Proline derivatives | 22.780 | L-Proline, N-valeryl-, heptadecyl ester | | $C_{27}H_{51}NO_3$ | 437 | 80 | Antiviral activity in plants (Abdelkhalek et al., 2022) and Antitumor agents (Bach & Takagi, 2013) |
| *Biemna fistulosa* | Diketopiperazines | 22.060 | 3,6-Diisopropylpiperazine-2,5-dione | | $C_{10}H_{18}N_2O_2$ | 198 | 77 | Biological activities, including enzyme inhibition and antimicrobial properties (Jia et al., 2022) |
| *Biemna fistulosa* | Diketopiperazines | 22.060 | Piperazine-3,5-dione, 1-tetradecanoyl | | $C_{18}H_{32}N_2O_3$ | 324 | 76 | |
| *Biemna fistulosa* | Fatty acids | 27.825 | Decanoic acid, 10-(2-hexylcyclopropyl) | | $C_{19}H_{36}O_2$ | 296 | 84 | Used in identifying bacteria and in research for studying its role in bacterial cell membrane protection and metabolism (Casillas-Vargas et al., 2021) |
| *Callispongia diffusa* | Quinolinedione derivative | 20.480 | 7-n-Pentadecylaminomethyl-6-hydroxy-5,8-quinolinedione | | $C_{25}H_{38}N_2O_3$ | 414 | 75 | Antibacterial; antifungal; antimalarial; anticancer and anti-inflammatory (Kadela-Tomanek et al., 2019) |
| *Callispongia diffusa* | Pyrazole derivative | 26.175 | 4H-Pyran-4-one, 2,2'-isopropylidene bis [3-methoxy-6-methyl] | | $C_{17}H_{20}O_6$ | 320 | 74 | Antibiotic, anti-inflammatory, antimalarial, antimicrobial, antiviral, anti-diabetic, anti-tumor, herbicidal, insecticidal, and analgesic activities (Almalki, 2023) |
| *Callispongia diffusa* | Lipophilic chemicals (phthalate acid esters) | 28.315 | Phthalic acid, 2-ethylhexyl tetradecyl ester | | $C_{30}H_{50}O_4$ | 474 | 87 | Antimicrobial, insecticidal, allelopathic and other biological activities (Huang et al., 2021) |
| *Callispongia diffusa* | Triglyceride (Triolein) | 27.830 | 9-Octadecenoic acid, 1,2,3-propanetriyl ester, (E, E, E) | | $C_{57}H_{104}O_6$ | 884 | 83 | Pharmaceuticals as a carrier or excipient in drug formulations; and an emollient and moisturizing therapy in skin care products; (N. Kumar & Sharma, 2023). |
| *Haliclona fascigera* | Sugar alcohols | 6.355 | Xylitol | | $C_5H_{12}O_5$ | 152 | 75 | Antibacterial, diabetes management, ear infection treatment, and gut health (Gasmi Benahmed et al., 2020); antimicrobial, dental and respiratory health (Salli et al., 2019); and anticancer, anti-inflammation and bone health (Ahuja et al., 2020); |
| *Haliclona fascigera* | Alkyl esters | 26.315 | n-Propyl 9-tetradecenoate | | $C_{17}H_{32}O_2$ | 268 | 72 | Antifungal activity against *Candida albicans* (Kadhim et al., 2016) |
| *Haliclona fascigera* | Esters Ergot alkaloids | 11.350 | Dodecanoic acid, 2-penten-1-yl ester | | $C_{17}H_{32}O_2$ | 268 | 78 | Anticancer, antioxidant, and antimicrobial activities (Hamouda et al., 2024) |

The methanolic extract of the marine sponge *B. fistulosa* (BLSi 007) revealed a total of 47 chemical compounds from the GC-MS analysis (Figure 6). These compounds were grouped into alkane derivatives (2.1%), amino acid derivatives (6.4%), cyclic alkanes (2.1%), cyclic amines (2.1%), cyclic dipeptides (2.1%), cyclic esters (2.1%), diketones (2.1%), diazabicyclo compounds (2.1%), ergot alkaloids (4.3%), ester derivatives (6.3%), esterified fatty acids (10.6%), esterified forms of ascorbic acids (vitamin C) (2.1%), fatty acids (2.1%), heterocyclic amines (10.6%), heterocyclic compounds (10.6%), hydrazones (2.1%), imidazoles (2.1%), ketones (4.3%), long-chain saturated fatty acids (4.3%), monounsaturated fatty acids (10.6%), organic nitrogenous compounds (2.1%), pyrazole derivatives (2.1%), triazole derivatives (2.1%), and unsaturated fatty acids (2.1%).

The GC-MS chromatogram data identified a total of 62 chemical compounds from the ethyl acetate extract of the sponge *C. diffusa* (BRMu 004) (Figure 7). The detected compounds were classified into alkenes (1.6%), amino acid derivatives (9.5%), amines (4.8%), aromatic hydrocarbons (4.8%), carboxylic acid anhydrides (1.6%), chiral amino alcohols (1.6%), chiral organic compounds (1.6%), esters (12.7%), ergot alkaloids (1.6%), fatty alcohols (1.6%), heterocyclic organic compounds (27.0%), ketals (1.6%), organic esters (15.9%), phthalic acid esters (7.9%), polycyclic hydrocarbons (1.6%), quinolinedione derivatives (1.6%), triglycerides (1.6%), and α-ketoglutaric acids (1.6%).

**Figure 7:**
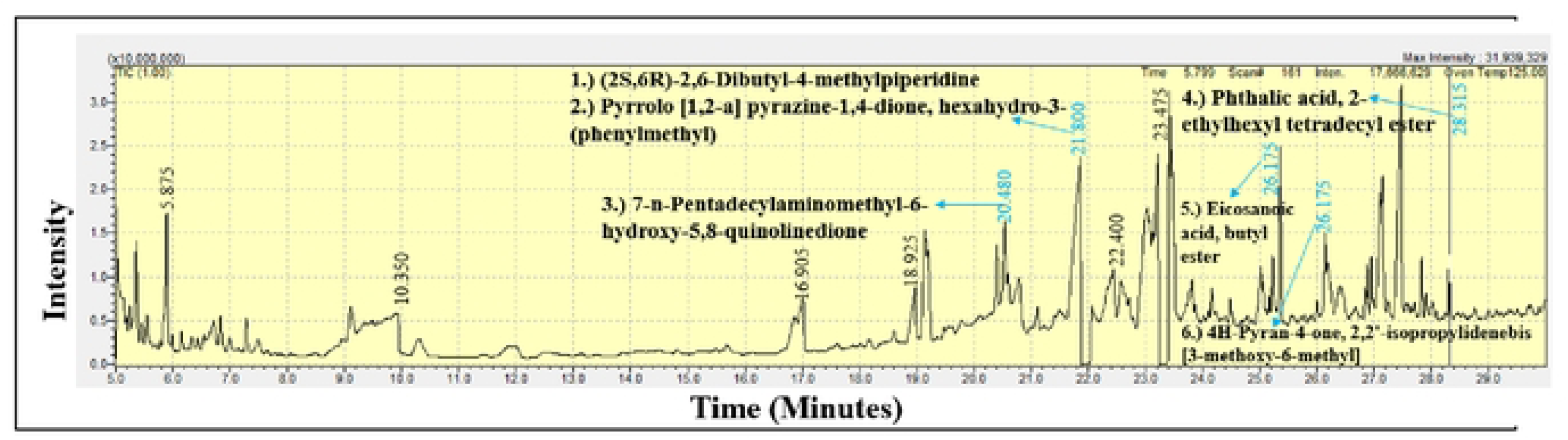
GC-MS chromatogram analysis of the ethyl acetate extract of *Callyspongia diffusa* (BRMu 004), identifying six secondary bioactive compounds.

The GC-MS analysis of the methanolic extract from the marine sponge *H. fascigera* (BLUCh 014) identified 37 chemical compounds (Figure 8). The chemical compound were categorized as acetylated amine alcohols (1.4%), amide compounds (2.7%), amide ester derivatives (13.5%), alkynes (2.7%), amino acid derivatives (2.7%), bicyclic chemical compounds (2.7%), cyclic alcohols (2.7%), cyclic alkanes (5.4%), cyclic amines (2.7%), epoxides (2.7%), esters (18.9%), heterocyclic amines (5.4%), heterocyclic boron-containing compounds (2.7%), heterocyclic compounds (10.8%), medium-chain fatty acids (2.7%), protected sugar derivatives (2.7%), pyrazole derivatives (2.7%), spiro compounds (2.7%), sugar alcohols (5.4%), synthetic compounds (2.7%), triazole derivatives (2.7%), and unsaturated carboxylic acids (2.7%).

**Figure 8:**
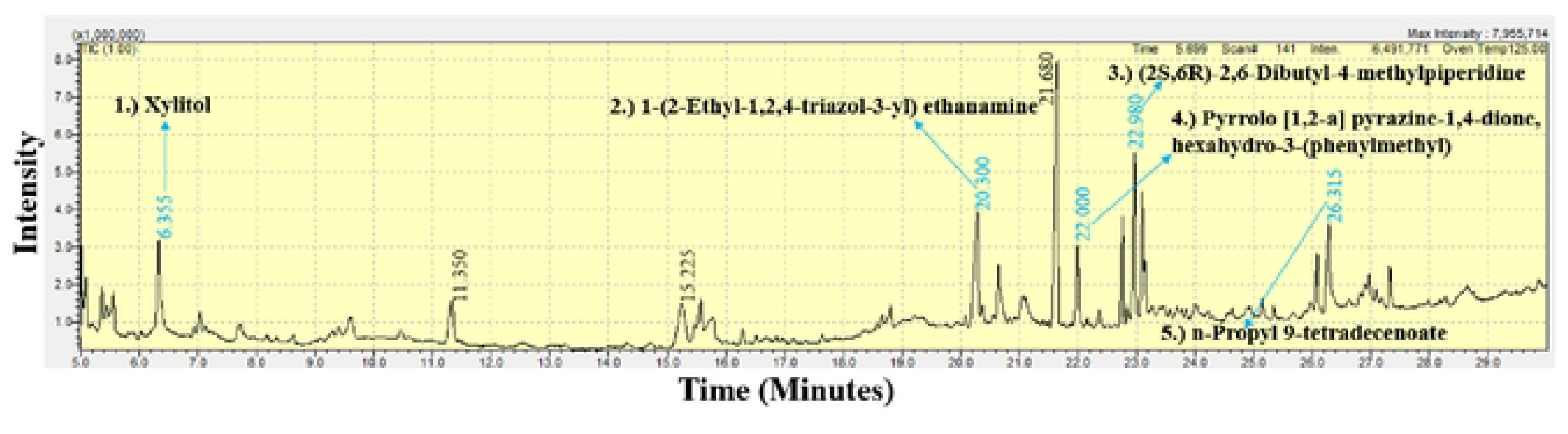
GC-MS chromatogram analysis of the methanolic extract of *Haliclona fascigera* (BLUCh 014), revealing five bioactive compounds.

In this study, GC-MS chromatogram analysis of the methanolic extract from *Biemna fistulosa* (BLSi 007) identified four potent bioactive compounds: (2S,6R)-2,6-dibutyl-4-methylpiperidine, pyrrolo [1,2-a] pyrazine-1,4-dione, hexahydro-3-(phenylmethyl), 1-(2-ethyl-1,2,4-triazol-3-yl) ethanamine, and 3,6-diisopropylpiperazin-2,5-dione (Figure 6 and Table 7).

Similarly, GC-MS analysis of the ethyl acetate extract from *Callyspongia diffusa* (BRMu 004) revealed six secondary bioactive compounds, including 7-n-pentadecylaminomethyl-6-hydroxy-5,8-quinolinedione, eicosanoic acid butyl ester, pyrrolo [1,2-a] pyrazine-1,4-dione, hexahydro-3-(phenylmethyl), 4H-pyran-4-one, 2,2’-isopropylidenebis [3-methoxy-6-methyl], (2S,6R)-2,6-dibutyl-4-methylpiperidine, and phthalic acid, 2-ethylhexyl tetradecyl ester (Figure 7 and Table 7).

The methanolic extract from *Haliclona fascigera* (BLUCh 014) exhibited five bioactive compounds: xylitol, 1-(2-ethyl-1,2,4-triazol-3-yl) ethanamine, pyrrolo [1,2-a] pyrazine-1,4-dione, hexahydro-3-(phenylmethyl), (2S,6R)-2,6-dibutyl-4-methylpiperidine, and n-propyl 9-tetradecenoate (Figure 8 and Table 6). Notably, the bioactive compounds (2S,6R)-2,6-dibutyl-4-methylpiperidine and pyrrolo [1,2-a] pyrazine-1,4-dione, hexahydro-3-(phenylmethyl) were consistently present in all three sponge extracts.

## DISCUSSION

In this study, the morphological identification of *Biemna fistulosa*, *Callyspongia diffusa*, and *Haliclona fascigera* revealed distinct skeletal structures and spicule compositions, aiding in their taxonomic classification. *B. fistulosa* exhibited a fibrous, encrusting growth pattern with curved diactinal styles and C-shaped sigma microscleres, while *C. diffusa* featured a spongin fiber reticulation with triangular spicule meshes, and *H. fascigera* is characterized by an isodictyal network of oxeas, raphides, and microstrongyles, commonly found in sandy lagoons and coral reef environments. A similar study used spicular analysis to investigate the sponge spicule assemblage, including *Biemna, Callyspongia*, and *Haliclona* species, in the lagoon reef of Bocas del Toro, Panama (Łukowiak, 2016). In Simeulue Island, Aceh Province, Indonesia conducted a morphological study was conducted on over twenty species of marine sponges. Their findings included notable species such as *Carteriospongia foliascens, Biemna fortis, Paratetilla aruensis, Oceanapia sp., Petrosia sp., Haliclona oculata,* and *Haliclona fascigera* (Setiawan et al., 2019) In addition, this study used a DNA barcoding, technique and identified three genera (*Biemna, Callyspongia*, and *Haliclona*) within the class Demospongiae. The complementary approaches significantly improved the precision of taxonomic classification for the marine sponges. The phylogenetic analysis clustered the sponge samples into three sub-clusters, with each representing a distinct genus. The specimen *Callyspongia* sp. BRMu004 (PQ329108) formed a distinct sub-cluster (supported by 100% bootstrap value) with members of the genus *Callyspongia* and had 100% sequence identity with *Callyspongia diffusa* (KX454494) (Figure 5; Table 1). On the other hand, *Haliclona* sp. BLUCh014 (PQ997929) clustered together with members of the genus *Haliclona* that had 99% sequence similarity. The specimen *Biemna* sp. BLSi007 (PQ997931) formed its sub-cluster (supported by 100% bootstrap value) with species from the genus *Biemna.* The formation of separate sub-clusters by the three sponge specimens on the phylogenetic tree indicated that they were representing distinct sponge species. A previous study examined species of shallow Hawaiian sponge fauna in the United States and identified *Biemna fistulosa* and *Callyspongia diffusa* using DNA barcoding (Núñez et al., 2017).

The GC-MS chromatogram data indicatted a total of 114 marine sponge chemical compounds from genera *Biemna, Callyspongia* and *Haliclona*. Notably, this study confirmed that 11.4% of the identified compounds had previously demonstrated antifungal, antibacterial, and antiviral bioactivity. A previous study demonstrated that *Callyspongia* species from the Red Sea exhibited antimicrobial properties, with bioactivity observed in the methanolic extract, its various fractions, and particularly in extracts purified from the dichloromethane fraction (Musa et al., 2022). Marine sponges belonging to the genus *Haliclona* are recognized for their capacity to biosynthesize a wide spectrum of secondary bioactive compounds such as sesquiterpenoid quinols, sterols, glycosphingolipids, and bioactive alkaloid compounds (Varijakzhan et al., 2021).

The findings in this study indicate that the ethyl acetate extracts of *C. diffusa* exhibited notable inhibitory activity against *P. aeruginosa*. The GC-MS analysis identified 62 chemical compounds in the ethyl acetate extract of *C. diffusa*, of which 9.7% have been previously reported to exhibit antimicrobial activity. These bioactive compounds include pyrrolo [1,2-a] pyrazine-1,4-dione, hexahydro-3-(phenylmethyl) (Al-Askar et al., 2024 and Kiran et al., 2018), (2S,6R)-2,6-Dibutyl-4-methylpiperidine (Ramya and Sethumadhavan, 2022), 7-n-Pentadecylaminomethyl-6-hydroxy-5,8-quinolinedione (Kadela et al., 2019), 4H-Pyran-4-one, 2,2’-isopropylidenebis [3-methoxy-6-methyl] (Almalki, 2023), phthalic acid, 2-ethylhexyl tetradecyl ester (Huang et al., 2021) and pyrrolo [1,2-a] pyrazine-1,4-dione; hexahydro-3-(2-methylpropyl)) (Askar et al., 2024 and Kiran et al., 2018). A similar study identified 212 bioactive compounds from the genus *Callyspongia,* and 109 molecules were reported to exhibit bioactivity (Sousa et al., 2021).

Notably, the methanolic extracts from the marine sponges *B. fistulosa* and *H. fascigera* show significant antibacterial activity against *E. coli.* Among the compounds identified in this study, 4.4% exhibited antibacterial activity, including pyrrolo[1,2-a] pyrazine-1,4-dione, hexahydro-3-(phenylmethyl) (Askar et al., 2024 and Kiran et al., 2018), 1-(2-ethyl-1,2,4-triazol-3-yl) ethanamine (Gupta et al., 2023), 7-n-pentadecylaminomethyl-6-hydroxy-5,8-quinolinedione (Kadela et al., 2019), 4H-pyran-4-one, 2,2’-isopropylidenebis [3-methoxy-6-methyl] (Almalki, 2023), and xylitol (Gasmi et al., 2020 and Salli et al., 2019). In a related study, extracts from *H. fascigera* sourced in Indonesia proved effective in inhibiting the growth of *S. aureus* and *E. coli* (Sugrani et al., 2019). Another study also revealed the antibacterial activities of crude extracts of *H. fascigera* against three shrimp pathogenic bacteria (Latifah et al., 2021). Elsewhere, Soulange et al. (2014) revealed that the organic sponge extracts exhibited greater antibacterial activity than the standard antibiotic against *S. aureus* and *E. coli*.

*Candida albicans* is the predominant causative agent of candidiasis, accounting for approximately 70% of fungal infections worldwide and contributing to the annual mortality of over 1.6 million people due to fungal diseases (Richardson, 2022). On one hand, *B. fistulosa* demonstrated more potent bioactive compounds capable of inhibiting *C. albicans* growth compared to *C. diffusa* and *H. fascigera*. Notably, 4.4% of the identified bioactive compounds were previously reported to possess antifungal properties, including pyrrolo [1,2-a] pyrazine-1,4-dione, hexahydro-3-(phenylmethyl) (Askar et al., 2024 and Kiran et al., 2018), pyrrolo[1,2-a] pyrazine-1,4-dione; hexahydro-3-(2-methylpropyl) (Al-Askar et al., 2024 and Kiran et al., 2018), 1-(2-ethyl-1,2,4-triazol-3-yl) ethanamine (Gupta et al., 2023), 7-n-pentadecylaminomethyl-6-hydroxy-5,8-quinolinedione (Kadela et al., 2019), and n-propyl 9-tetradecenoate (Kadhim et al., 2016). Xestodecalactones B compounds isolated from Xestospongia *exigua* in the Bali Sea demonstrated antifungal activity against *C. albicans* (Varijakzhan et al., 2021). Similarly, nortetillapyrone, a tetrahydrofurylhydroxypyran-2-one derived from *Haliclona cymaeformis*, exhibited antifungal efficacy against various fungal pathogens with distinct MIC values (Ngece et al., 2024). Furthermore, polyketide compounds such as woodylides A and C from *Plakortis simplex*, as well as theonellamide G, swinholide I, and hurghadolide A from *Theonella swinhoei*, along with tetramic acid glycosides aurantosides G and I, demonstrated significant antifungal activity, underscoring the potential of marine sponges as reservoirs of antifungal compounds (Yu et al., 2012).

The decanoic acid, 10-(2-hexylcyclopropyl), a fatty acid obtained from the methanolic extract of *B. fistulosa*, has been used in identifying bacteria and in research for studying its role in bacterial cell membrane protection and metabolism (Casillas et al., 2021). Furthermore, 9-octadecenoic acid, 1,2,3-propanetriyl ester E, E, E, a triolein compound, was extracted from the ethyl acetate extract of *C. diffusa*. This compound holds pharmaceutical significance as a carrier or excipient in drug formulations and is also employed as an emollient and moisturizing agent in dermatological products (Kumar and Sharma, 2023). In a previous study, the bioactive compound 9-Octadecenoic acid, 1,2,3-propanetriyl ester (E, E, E) was discovered in *Cassia angustifolia* and demonstrated significant antimicrobial activity (Marzoqi et al., 2016).

In our study, the methanolic extract of *H. fascigera* exhibited a lower minimum inhibitory concentration (MIC), categorizing it as a potent inhibitor with enhanced efficacy in suppressing the growth of *Escherichia coli* compared to the standard drug. Moreover, the extract demonstrated significant bactericidal activity, with a minimum bactericidal concentration (MBC) of 1.25 mg/mL, which was substantially lower than the MBC values recorded for *C. diffusa* and *B. fistulosa*. Additionally, this research established that the ethyl acetate extract derived from *C. diffusa* demonstrated bactericidal activity against *Pseudomonas aeruginosa*, with an MBC value of 2.5 mg/mL. Marine sponge extracts from a study of the waters of Mauritius exhibited low MBC values, indicating that these extracts could serve as a potential approach to traditional bacterial infection management strategies (Beesoo et al., 2017). In a similar study, *H. fascigera*, collected from Bidong Island, Malaysia, showed positive results in all in vitro biological studies and exhibited higher antibacterial activities compared to other sponges in the study (Jamaludin et al., 2023). The research also revealed that the butanol fraction of *Biemna* sp. exhibited significant antibacterial activity, with an MIC value of 0.091 mg/ml against *Escherichia coli* (Soulange et al., 2014).

The findings of this study indicate that the methanolic extract of *H. fascigera* exhibited the most potent fungicidal activity, with a minimum fungicidal concentration (MFC) of 2.5 mg/mL. This suggests that the extracts of *H. fascigera* are more effective at eliminating *C. albicans* at lower concentrations compared to the other extracts evaluated. Furthermore, research conducted in Biak, Indonesia, demonstrated that the ethyl acetate extracts of *Fascaplysinopsis* sp. and *Haliclona* sp. possess notable antifungal activity against *C. albicans* (Kasitowati et al., 2019). Additionally, investigations on marine sponges from the Ratnagiri coast of India reported moderate antifungal activity against *C. albicans* (Kumar, 2022).

## CONCLUSION AND RECOMMENDATIONS

This research study underscores the significance of marine sponges as a promising reservoir of antimicrobial agents. Extracts derived from *B. fistulosa*, *C. diffusa*, and *H. fascigera* exhibited pronounced antibacterial activity compared to the positive control. Furthermore, the methanolic extract of *H. fascigera* demonstrated the most potent antifungal activity among the marine sponge extracts evaluated. The findings of this study suggest that marine sponges from Kenyan waters possess notable therapeutic potential, presenting valuable lead compounds for drug discovery and development. More studies should focus on the mechanisms of action and toxicity of pure leads isolated from sponges at the molecular level, which is important to give direction for lead improvement and further drug development.

## AUTHORS’ CONTRIBUTION

Conceptualization, T.N.W., H.M.M., J.K.N, C.M.A, and C.M.K; writing— original draft preparation, T.N.W.; writing—review and editing, T.N.W., H.M.M., J.K.N, C.M.A and C.M.K. All authors have read and agreed to publish the manuscript.

## DATA AVAILABILITY STATEMENT

The data presented in this study are available upon request from the corresponding authors.

## ACKNOWLEDGEMENTS

We are profoundly grateful to the Kenya Marine and Fisheries Research Institute (KMFRI) and the Technical University of Mombasa (TUM) for their unwavering support. Their outstanding laboratory facilities were instrumental in the successful completion of this research, enabling us to achieve our objectives effectively.

